# Hepatocyte expression of fetal insulin receptor isoform contributes to the promotion of liver cancer through non-cell autonomous mechanisms

**DOI:** 10.1101/2025.06.02.655477

**Authors:** Fanny Léandre, Akila Iddir, Cécile Godard, Isabelle Lagoutte, Anthony Caldiero, Soumeya Souid, Sandra Pinto, Jérémy Augustin, Vanessa Bou Malham, Alexandre Alves, Rowan Aubry, Sandrine Imbeaud, Jessica Zucman-Rossi, Sabine Colnot, Angélique Gougelet, Christèle Desbois-Mouthon

**Affiliations:** INSERM, Université Paris Cité, Sorbonne Université, Centre de Recherche des Cordeliers, F-75006 Paris, France; INSERM, CNRS, Université Paris Cité, Institut Cochin, Plateforme Imagerie du vivant, F-75014 Paris, France; INSERM, Université Paris Est Créteil, IMRB, F-94010, Créteil, France; Assistance Publique-Hôpitaux de Paris, Hôpital universitaire Henri Mondor-Albert Chenevier, Département d’anatomopathologie, F-94010, Créteil, France

**Keywords:** Hepatocellular carcinoma, hepatoblastoma, liver, alternative splicing, insulin receptor, β-catenin, apoptosis, inflammation, CRISPR/Cas9, dependence receptor

## Abstract

The insulin receptor (INSR) exists in two isoforms, INSR-A and INSR-B, resulting from alternative splicing of the *INSR* pre-mRNA. INSR-B mediates the metabolic and mitogenic effects of insulin in the adult liver, while INSR-A is expressed during development. Recently, INSR-A has been detected in pathological murine and human livers. Here, we develop an *in vivo* CRISPR/Cas9 strategy to assess the impact of INSR-A on mouse liver homeostasis and susceptibility to carcinogenesis. We find that INSR-A expression in hepatocytes leads to the spontaneous development of liver tumours and also increases tumour initiation in a context of β-catenin-driven liver carcinogenesis. Mechanistically, this is attributed to the higher intrinsic capacity of INSR-A expressing hepatocytes to enter apoptosis, rendering the microenvironment more inflammatory, thus making way for the proliferation of preneoplastic cells. Collectively, our data highlight a novel function for INSR-A in promoting liver cancer *via* non-cell autonomous mechanisms.

## Introduction

In the liver, the insulin receptor (INSR) mediates some, but not all, of insulin’s effects on carbohydrate and lipid homeostasis.^1^ Thus, the constitutive or inducible loss of hepatic INSR leads to glucose intolerance combined with hyperinsulinaemia in liver-specific INSR knockout (LIRKO) mice.^2–4^ Evidence also suggests that the INSR plays a contributory role to the pathogenesis of liver diseases. Mice with INSR hepatic loss are protected against the accumulation of excess hepatic triglycerides and do not develop fatty liver disease in response to a high-fat diet.^5,6^ Recently, an atlas of the liver kinome identified the INSR as one of the most hyperactivated receptor in human and rodent fibrotic livers.^7^ Moreover, membranous INSR overexpression has been associated with worse survival in patients with primary liver cancer.^8^

The molecular mechanisms whereby the INSR intrinsic tyrosine kinase (TK) is activated upon insulin binding have been well characterized in insulin target cells. In response to insulin, INSR autophosphorylates and initiates a cascade of phosphorylation events leading to the rapid activation or inhibition of intracellular enzymes and the regulation of gene expression involved in the stimulation of glycogen synthesis, lipogenesis, and lipoprotein synthesis and the suppression of gluconeogenesis, glycogenolysis and VLDL secretion.^9^

The existence of two INSR isoforms adds a level of complexity to INSR signalling. INSR is present as two isoforms, INSR-A and INSR-B, which result from the alternative splicing of exon 11 from *INSR* pre-mRNA.^10^ INSR-B is the long isoform (exon 11 inclusion) expressed in adult hepatocytes which relays insulin actions.^11^ The short INSR-A isoform (exon 11 skipping), is often referred to as oncofetal INSR, as it is expressed during tissue development and re-expressed in various cancer tissues and cell lines.^12^ We found that INSR-A is expressed in human liver cancer cell lines and tumours at the expense of INSR-B due to deregulation of *INSR* pre-mRNA splicing.^13,14^ In addition, mRNA encoding INSR-A has been detected in murine and human pathological but non-tumourous livers.^15–17^

The description of the functions assigned to INSR-A was carried out mainly *in vitro*, using stable overexpression cellular systems. While INSR-B binds insulin with a high affinity, INSR-A binds not only insulin but also insulin-like growth factor-II (IGF-II) and pro-insulin.^18–20^ It has been reported that, after stimulation of mouse fibroblasts with insulin, overexpressed INSR-A drives pathways involved in cancer, stemness and interferon signaling, while overexpressed INSR-B induces pathways involved mainly in metabolic or tumour suppressive functions.^21^ In human liver cancer cell lines, the overexpression of INSR-A promotes an aggressive phenotype associated with increased migratory and invasive properties and stem/progenitor-cell features.^14^

Hepatocellular carcinoma (HCC) is the most prevalent primary liver cancer, ranking as the sixth most common cancer and the third-leading cause of cancer-related mortality worldwide.^22^ HCC occurs in 80%–90% of patients with underlying cirrhosis due to viral, alcoholic or metabolic chronic liver disease. The mutational spectrum in HCC has been well characterized, and indicates that 30-40% of tumours have *CTNNB1* mutations which activate the β-catenin pathway and associate with G5/G6 subgroups in the Boyault *et al.* molecular classification.^23,24^ *CTNNB1*-mutated HCC demonstrate unique gene expression patterns and pathological features.^25,26^ They are usually low-proliferating tumours with a high degree of differentiation, prominent cholestasis, a peculiar metabolism and the absence of immune infiltration. They show robust immunohistochemical expression of glutamine synthetase (GS), a direct target of the β-catenin pathway.

In the current study, we investigated the functions that can be assigned to INSR-A *in vivo* in normal adult liver or adult liver under oncogenic stress. To this end, we used an *in vivo* CRISPR/Cas9 technology to generate a mouse model whose hepatocytes were genetically edited to express INSR-A. We showed that INSR-A expression has no impact on liver homeostasis in the short term, but that it is associated with spontaneous tumour development in the long term and also promotes liver carcinogenesis in a context of β-catenin oncogenic activation. Unexpectedly, we found that the vast majority of tumours derives from non-edited hepatocytes. We found that expression of INSR-A provoked an inflammatory response associated with increased apoptosis of edited hepatocytes and compensatory proliferation of β-catenin-activated hepatocytes in the liver parenchyma. Our results highlight the impact of INSR-A on liver homeostasis and the promotion of hepatocyte transformation, through non-cell autonomous effects which could be attributed to the higher intrinsic susceptibility of INSR-A-expressing hepatocytes to die.

## Material and methods

### Mouse models and procedures

Mice were housed on a 12/12-h light/dark cycle at 22°C-24°C with *ad libitum* access to food (Safe A03, SAFE® Complete Care Competence) and water. Except when specified, 2-3-month old male mice were used throughout the study. The generation of APC^ΔHep^ and βCAT^ΔEx3^ mice has been described previously.^27,28^ Tumour development was followed by 2D-ultrasounds, as previously described.^29^ Retro-orbital injections and ultrasonography were performed under anesthesia with isoflurane inhalation. For terminal tissue and blood collection, mice were anesthetized with ketamine (80 mg/kg) and xylazine (5 mg/kg). Mice were fasted 5-6 h before sacrifice. All animal experiments were carried out in accordance with the French government regulations and with the approval of two ethical committees (Charles Darwin, Sorbonne Université, APAFIS#16420, APAFIS#45107 and CEEA34, Université de Paris, APAFIS#14472)

### Glucose tolerance test (GTT), insulin tolerance test (ITT) and *in vivo* insulin signalling

After a 6 h fast, mice were intraperitoneally injected with glucose (1.5 g/kg body weight) and blood samples were taken at 0, 15, 30, 60, and 120 min from the tail vein. Glucose levels were measured using an Accu-Chek Performa glucometer (Roche). For insulin tolerance test (ITT), 5 h fasted mice were injected intraperitoneally with human insulin (Umuline® NPH, 1 IU/kg body weight, Lilly France). Blood glucose levels were measured as previously. Insulin levels were measured in plasmas using the ultrasensitive mouse insulin ELISA kit, 2^nd^ Gen (Crystal Chem). For *in vivo* insulin signalling studies, fasted mice were injected intraperitoneally with human insulin (1 U/kg body weight). Saline-injected animals served as controls. After 10 min, livers were removed and immediately frozen in liquid nitrogen.

### Liver function

Alanine transaminase (ALT) and aspartate transaminase (AST) activity was measured in mouse plasma using dedicated kits (ThermoFisher Scientific) on an automated biochemistry analyzer (Indiko plus, ThermoFisher Scientific).

### Cell culture, transfection and treatments

Mouse hepatocytes were isolated from perfused livers, as previously described.^29,30^ Murine Hepa1-6 hepatoma cell line (ATCC® CRL-1830^TM^) was cultured in Dulbecco’s Modified Eagle’s Medium (DMEM) supplemented with 10% fetal calf serum (FCS) and antibiotics. Human PLC/PRF5 cells (gift from Dr Christine Perret, Institut Cochin, Paris) were cultured in MEM with 10% FCS, pyruvate, and non-essential amino acids. Cells were seeded in six-well plates and transfected using 2.5 μg of total plasmid DNA mixed with Lipofectamine 2000, according to manufacturer’s instructions (ThermoFisher Scientific). Primary hepatocytes were incubated with either staurosporine (10 μM, Medchem Express) or a combination of staurosporine (3.33 μM) and TNF-α (7 ng/ml, Biolegend).

### Design of small guide (sg) RNAs for *INSR/Insr* gene editing by *Staphylococcus aureus* Cas9 (saCas9) nuclease

Several sgRNAs of 21 nucleotides located in introns 10 and 11 from human *INSR* and murine *Insr* were selected according to their GC content (20:80%) and low potential off-target sites using the web-based tool Cas-designer (http://www.rgenome.net/cas-designer/).^31^ A pair of complementary oligonucleotides with 5’-CACC and 5’-AAAC overhangs were synthesized for each human and murine targeted site, treated with T4 polynucleotide kinase, annealed, and cloned into the BsaI site of the viral vectors pX601 (#61591, saCas9 cDNA under the cytomegalovirus (CMV) promoter) and pX602 (#61593, saCas9 cDNA under the thyroxine-binding globulin (TBG) promoter)^32^ (Addgene, gifts from Pr Feng Zhang) (**Figure S1A**). To test *INSR*/*Insr* gene editing efficiency, recombinant pX601 vectors were individually transfected into human PLC/PRF5 and murine Hepa1-6 cell lines. The spectrum and frequency of mutations generated were evaluated in genomic DNA after PCR amplification of the region of interest, purification, sequencing and tracking of INDELS by decomposition (TIDE method). The pairs of constructs encoding the most effective sgRNAs (**Table S1, Figure S1B**) were then transiently co-transfected in Hepa1-6, primary murine hepatocytes and PLC/PRF5 cells. For murine cells, the conditions under which controls were performed used the same vector backbone in which a sgRNA targeting the neutral locus *Rosa26* was inserted.^28^ For human cells, non-edited clones served as controls.

### Genomic DNA isolation and PCR amplification

Tissue and cell samples were lysed overnight at 56°C in 50 mM Tris-HCl pH 8.0, 50 mM EDTA, 100 mM NaCl, and 1% SDS supplemented with 0.8 mg/ml proteinase K (Promega). Genomic DNA was extracted using a conventional procedure with ultrapure phenol:chloroform:isoamyl alcohol (25:24:1, v/v, ThermoFisher Scientific). Genotyping PCR was performed using 400 ng DNA and Taq HS (Ozyme) according to the manufacturer’s protocol and was analyzed by electrophoresis on a 2% agarose gel (Invitrogen^TM^ E-Gel precast agarose gels with SYBR Safe DNA gel stain, ThermoFisher Scientific). PCR primer pairs are given in **Table S2**. The deletion of exon 11 and its adjacent intronic regions in *Insr* and *INSR* resulted in the appearance of an additional 423-bp fragment (mouse wild-type fragment: 1059 bp) and a 360-bp fragment (human wild-type fragment: 711 bp), respectively (**Figure S1B**).

### Total RNA Isolation, reverse-transcription and quantitative real-time PCR (RT-QPCR)

Total RNA was extracted from liver tissues using Trizol reagent (ThermoFisher Scientific) according to the manufacturer’s protocol. Total RNA was extracted from cell cultures using Nucleospin RNA kit (Macherey-Nagel SARL). cDNA was synthesized using Maxima reverse transcriptase (ThermoFisher Scientific). RT-QPCR was performed with specific primers using BlasTaq^TM^ 2X PCR master mix (abm) or TaqMan^TM^ gene expression assays (Applied Biosystems). The expression levels of each target gene were normalized to those of *HPRT*/*Hprt* and expressed as the relative quantity using the formula 2^-ΔΔCt^. PCR primer pairs and TaqMan gene expression assays are listed in **Tables S2** and **S3**.

### Gene expression microarray

Total RNA was extracted using Trizol (ThermoFischer Scientific) from 3-4 tumours of each group. RNA integrity was assessed using the Agilent 2100 Bioanalyzer (Agilent Technologies). Total RNA (300 ng) was reverse transcribed, amplified and labelled using the GeneChip^TM^ WT PLUS Reagent Kit (Affymetrix). Each cDNA sample was hybridized to GeneChip^®^ Clariom S Mouse (Life Technologies). After overnight hybridization at 45°C, chips were washed on the fluidic station FS450 following specific protocols (Affymetrix) and scanned using the GeneChip^TM^ Scanner 3000 7G (ThermoFischer Scientific). The scanned images were analyzed with Expression Console software (Thermo Fisher Scientific) to obtain raw data (cel files) and metrics for quality controls. Raw data were normalized using the robust multichip algorithm (RMA) and TAC4.0 software (Thermo Fisher Scientific). Datasets are available at GSEXXX. RNAseq data comparing APC^Δhep^ livers to APC^WT^ livers has been published previously (project: PRJNA150641 in European nucleotide archive (ENA).^28,29,33^

### Characterization of PLC/PRF5 cells with CRISPR/Cas9-induced *INSR* gene edition

PLC/PRF5 cells were transfected with plasmids pX601/CMV encoding each sgRNA against *INSR* locus, together with a puromycin resistance plasmid. 24h later, cells were treated with puromycin (4 μg/ml) for a 48 h period. A clone edited in one *INSR* allele (#1F7) was obtained by limiting dilution and amplified. Unedited clones were used as controls (#1A9, #2H6). Clones were tested for their invasive capacity using Matrigel®-coated Transwells® (Corning) and were also tested for their ability to form spheroids in suspension as previously reported.^14^

### Production and injection of recombinant AAV8-saCas9-sgRNA viral particles

Viral vectors containing sgRNA against *Insr* or *Rosa26* were amplified in the Stbl3 bacterial strain, purified using NucleoBond Xtra Maxi Plus EF maxi kit (Macherey-Nagel) and then sent to the vector production center (INSERM UMR1089, Université de Nantes, France) for recombinant AAV8 production. Unless otherwise specified, a total of 4×10^11^ genomic copies (2×10^11^ for each AAV8-saCas9-sg*Insr* or 4×10^11^ for AAV8-saCas9-sg*Rosa*) were diluted in 150 μl of sterile physiological serum and retro-orbitally injected into the mice.

### Liver histology, immunohistochemistry (IHC), steatosis and TUNEL assay

Hematoxylin-Eosin (H&E) staining, IHC and TUNEL assay were carried out on formalin-fixed paraffin-embedded (FFPE) liver tissue sections. For the assessment of liver steatosis, 10-μm liver frozen sections were fixed in 10% formalin for 10 min, rinsed in 60% isopropyl alcohol, and then incubated in Oil Red O solution (Sigma) for 15 min. IHC staining for myeloperoxidase (#22225-1-AP, Proteintech), glutamine synthetase (GS) (#610518, BD Transduction Laboratories), Ki67 (#MA5-14520, SP6, ThermoFisher Scientific), cleaved caspase-3 (#9661, D175) and F4/80 (#70076, D2S9R, Cell Signaling) was performed according to the provider instructions. Detection of *in situ* apoptosis was assessed by terminal deoxynucleotidyl transferase dUTP nick-end labeling (TUNEL) assay, performed according to the manufacturer’s instructions (Click-iT™ TUNEL Alexa Fluor™ 647, ThermoFisher Scientific). Photographs of whole slides were taken using the AxioScan Z1 scanning system (Zeiss). The number of stained cells per field was determined using ImageJ (Version 1.52a, National Institute of Health) or Qupath ^34^ software.

### *In situ* hybridization

4-μm FFPE sections of mouse livers were used for *in situ* hybridization using Base Scope™ paired double-Z oligonucleotide probes (ACDBio). The Insr-A probe (BA-Mm-Insr-tv1-E10E11) targeting 2700-2735 positions of transcript NM_010568.3 was visualized in red. Negative (bacterial *dapb*) and positive (mouse *Ppib* and *Polr2a*) control probes were tested in parallel. Negative controls showed a negligible background. Photographs of whole slides were taken using the AxioScan Z1 scanning system (Zeiss).

### Western blot (WB) analysis

For protein extraction, whole-liver tissue and primary hepatocytes were homogenized in radioimmunoprecipitation assay buffer together with cocktail of protease and phosphatase inhibitors (Sigma-Aldrich). 30-100 μg of protein lysates were resolved on 10-15% polyacrylamide gels, transferred to nitrocellulose membranes, and blocked with 5% BSA or milk for 1 h before incubation with primary antibodies overnight at 4°C. Membranes were incubated with secondary antibodies (Cell Signaling Technology), revealed with Clarity^TM^ Western ECL substrate (Bio-Rad) and protein bands visualized with the iBright^TM^ CL1500 imaging system (ThermoFisher Scientific). Signal intensity was quantified using Image J. Antibodies specific to phospho-INSR β-subunit (#03024, 19H7), INSR β-subunit (#3025, 4B8), phospho-AKT (#4060, D9E), AKT (#4691, pan C67E7), phospho-PRAS40 (#13175, D4D2), PARP (#9542), cleaved caspase-3 (#9661, D175) (Cell Signaling Technology) and IGF-II were used (#Ab9572, Abcam). Protein loading was controlled with a β-actin (#A5441, Sigma) antibody.

### Cytokine/chemokine array

Proteins were extracted from liver tissues in PBS supplemented with protease inhibitors (10 μg/ml aprotinin, 10 μg/ml leupeptin, and 10 μg/mL pepstatin A) and 1% Triton® X-100 (Sigma). 200 μg of protein was analyzed using the mouse XL cytokine array (R&D Systems) according to the manufacturer’s instructions. The arrays were revealed and quantified as performed in WB analysis.

### Human RNA-seq datasets

We used bulk RNA-seq datasets previously published in European Genome-phenome archive (EGA) for extracting information relative to INSR isoform mRNA expression. These datasets include 48 pediatric non-tumour livers (EGAS00001006692, EGAS00001005108), 73 adult non-tumour livers (EGAS00001004629, EGAS00001003837, EGAS00001007694, EGAS00001002091, EGAS00001007957, EGAS00001005986, EGAS00001003685), 98 hepatoblastoma (EGAS00001005108, EGAS00001006692) and 427 HCC (EGAS00001005108, EGAS00001003837, EGAS00001002879, EGAS00001004629, EGAS00001007694, EGAS00001007957, EGAS00001001284, EGAS00001008084, EGAS00001003310, EGAS00001005986). Gene-expression-based classifications of HCC and HB were done as previously described.^35^

### Statistical analysis

Data are presented as median and interquartile range (IQR; 25%–75%). The Mann-Whitney U test was used to compare differences between two unpaired groups. A Gehan-Breslow-Wilcoxon test was used to compare the cumulative number of tumour-free mice. Spearman’s rank correlation coefficient was used to evaluate the correlation. Statistical analysis was performed using GraphPad Prism 6.0, with the significance set at *P* < 0.05.

## Results

### The CRISPR/Cas9 strategy is effective in promoting the expression of INSR-A at the expense of INSR-B in hepatocytes

To analyze the influence of INSR-A on hepatic homeostasis and carcinogenesis in adult mice, we developed an *in vivo* CRISPR/Cas9 gene-editing system to promote excision of *Insr* exon 11 and thus to favour the expression of *Insr-a* mRNA to the detriment of *Insr-b* mRNA in hepatocytes. As a proof of concept and before starting *in vivo* experiments, we first validated the strategy *in vitro* using human hepatic PLC/PRF5 cells transfected with plasmids encoding two sgINSR and saCas9 under *CMV* promoter (**Figures S1** and **S2**). The heterozygous *INSR*-edited clone 1F7 exhibited exon 11 excision and increased *INSR-A* mRNA level together with decreased *INSR-B* mRNA levels, as compared to unedited control clones 1A9 and 2H6 (**Figures S2A-S2C**). Clone 1F7 also showed increased invasive properties (**Figure S2D**), and a higher propensity to form spheroids in non-adhesive conditions, as compared to controls (**Figure S2E**). These results are in full agreement with our previous study,^14^ which showed that the ectopic overexpression of INSR-A in PLC/PRF5 cells enhances cell invasion and progenitor cell features. The CRISPR/Cas9 strategy we had developed was therefore effective in replacing the endogenous INSR-B with an INSR-A capable of mediating aggressive features in human liver cells.

Based on these results, we next developed and validated the CRISPR/Cas9 tools to target *Mus musculus Insr* using the murine cell line Hepa1.6 (**Figure S3A**). Recombinant hepatotropic AAV8 expressing sgInsr or sgRosa together with saCas9 under *TBG* promoter were then produced. A dose-effect study showed that, after a duration of one month, the injection of 2.10^11^ viral genomes (vg) efficiently induced the excision of *Insr* exon 11 and flanking regions in genomic DNA from whole liver (median: 33.50%; IQR: 31.25-40.50, n=16) but not from other organs (**Figures S3B-S3C**). Accordingly, this dose led to a significant induction of *Insr-a* mRNA (median: 4.5-fold; IQR: 3.0-6.4) together with a significant downregulation of *Insr-b* mRNA (median: 0.6-fold; IQR: 0.5-0.8) in the whole liver while total *Insr* mRNA transcripts remained unchanged (**Figure 1A**). The induction fold for *Insr-a* mRNA was higher in freshly isolated hepatocytes (median: 15.6-fold; IQR: 10.8-19.1) (**Figure 1B**). Since there is no antibody specific to INSR-A, *in situ* hybridization was used to detect *Insr-a* transcript on liver tissue. Thus, upregulation of *Insr-a* mRNA expression was shown to be hepatocyte-specific at single-cell level in sgInsr liver (**Figure S3D**). All these data confirm that CRISPR-Cas9-mediated gene editing can be used to mimic alternative splicing of *Insr* pre-mRNA in the hepatocytes.

**Figure 1:**
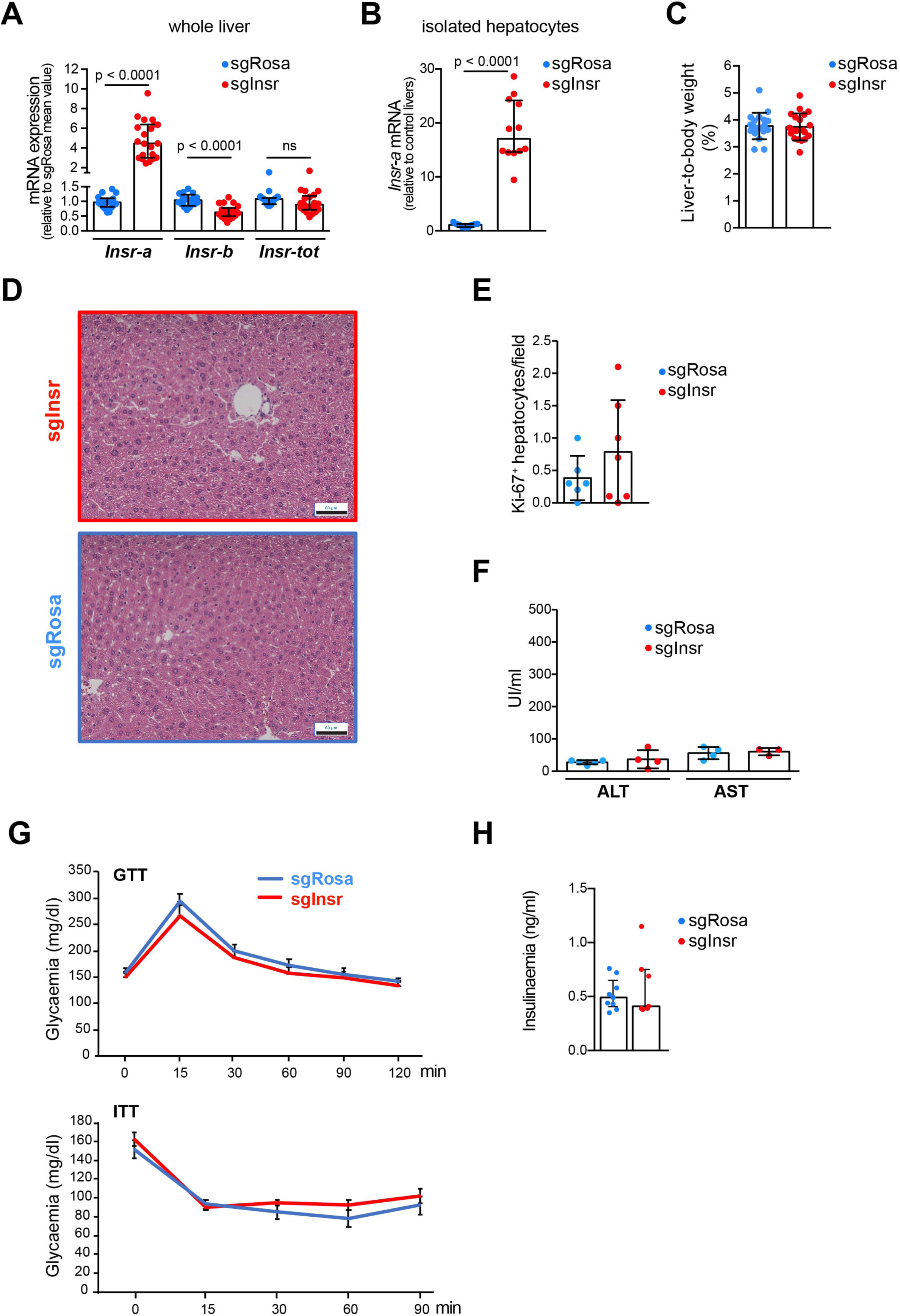
Expression of INSR-A in hepatocytes has no impact on liver homeostasis in the short term. Mice were injected with recombinant AAV8 encoding saCas9 and sgRNA against *Rosa* (sgRosa) or *Insr* (sgInsr) locus. Analyses were performed between 6-11 weeks after injection. **A.** Expression of total *Insr* and specific *Insr*-a/*Insr-b* transcripts in whole livers evaluated by RT-qPCR (n=20). **B.** Expression of *Insr*-a transcript by RT-qPCR in freshly isolated hepatocytes (n=12). **C.** Liver-to-body weight ratio (n=16). **D.** Representative images of H&E staining. Scale bar: 60 μm. **E.** Quantification of Ki67-positive proliferating hepatocytes by immunohistochemistry (n=6-7). **F.** Plasma transaminase levels (alanine transaminase (ALT), aspartate transaminase (AST)) (n=3-5). **G.** Glucose tolerance test (GTT, *upper panel*) and insulin tolerance test (ITT, *lower panel*) performed on 5-6 h fasted mice (n=8 for each time point). **H.** Insulinaemia measured on 5-6 h fasted mice (n=8-9). Data are median ± IQR.

### Expression of INSR-A in hepatocytes has no impact on liver homeostasis in the short term

We examined whether hepatocyte expression of INSR-A impacted liver homeostasis in the short term. Six weeks after AAV8-sgInsr injection, no changes in liver-to-body weight ratio (**Figure 1C**), liver histology (**Figure 1D**), hepatocyte proliferation (**Figure 1E**), or liver function (**Figure 1F**) were observed compared with control AAV8-sgRosa injected mice. ALT and AST plasma levels remained unmodified 11 weeks after injection (**Figure S4A**). Glucose tolerance tests (GTT) and insulin tolerance tests (ITT) did not reveal any disturbance in glucose disposal and insulin sensitivity in AAV8-sgInsr injected mice, respectively (**Figure 1G**). Both groups of mice had similar insulin levels after short-term fasting (**Figure 1H**). Finally, no changes in INSR, AKT and PRAS phosphorylation were observed in the livers of AAV8-sgInsr mice after insulin injection (**Figure S4B**).

### Expression of INSR-A in hepatocytes leads to tumour development in the long term

Liver ultrasound was then used to monitor tumor development in male mice injected with AAV8-sgInsr (n=10) and AAV8-sgRosa (n=8) for up to 13 months (**Figure 2A**). Weight gain was similar between the two groups over that period (**Figure S5A**). One AAV8-sgRosa mouse had lymphoma at the time of sacrifice and was excluded from the analysis. Another died 11 months after injection, but was tumour-free. Three AAV8-sgInsr mice developed one tumour ∼200-300 days after injection while no tumour was detected in the remaining AAV8-sgRosa mice (**Figures 2B** and **2C**). Surprisingly, genotyping PCR revealed that no tumour was edited at the *Insr* locus, in contrast to AAV8-sgInsr nontumourous livers (NTL) (**Figure 2D**), which also showed increased expression of *Insr-a* mRNA compared to AAV8-sgRosa NTL (**Figure 2E**). The three tumours were proliferative (**Figure 2F**), did not express GS (**Figure 2G**) and two of them overexpressed *Afp* and *Gpc3* mRNA (**Figure 2H**). One tumour was identified as well-differentiated steatotic HCC and two tumours as hepatocellular adenomas, but the differential diagnosis was difficult even after reticulin staining (data not shown). Histological analysis of steatosis in NTL revealed equivalent levels in both groups (**Figure 2I**) associated, however, with higher expression of certain mRNAs encoding the core enzymes of *de novo* lipogenesis in AAV8-sgInsr NTL (**Figure S5B**). Molecular analysis of fibrosis showed no difference between the two groups (**Figure S5C**). Fasting insulinaemia at sacrifice was similar between the two groups (**Figure 2J**). Finally, AAV8-sgInsr were also injected into female mice (n=10), none of which developed tumours after 13 months. Overall, these data suggest that INSR-A expression in hepatocytes has no impact on liver homeostasis in the short term, but leads to the spontaneous development of benign and malignant liver tumours in the long term in males, with a penetrance of 30%, in a context of liver steatosis. All tumours retained the wild-type *Insr* gene, indicating that *Insr*-edited hepatocytes are not the origin of these tumours, but may participate in their initiation via extrinsic signals.

**Figure 2:**
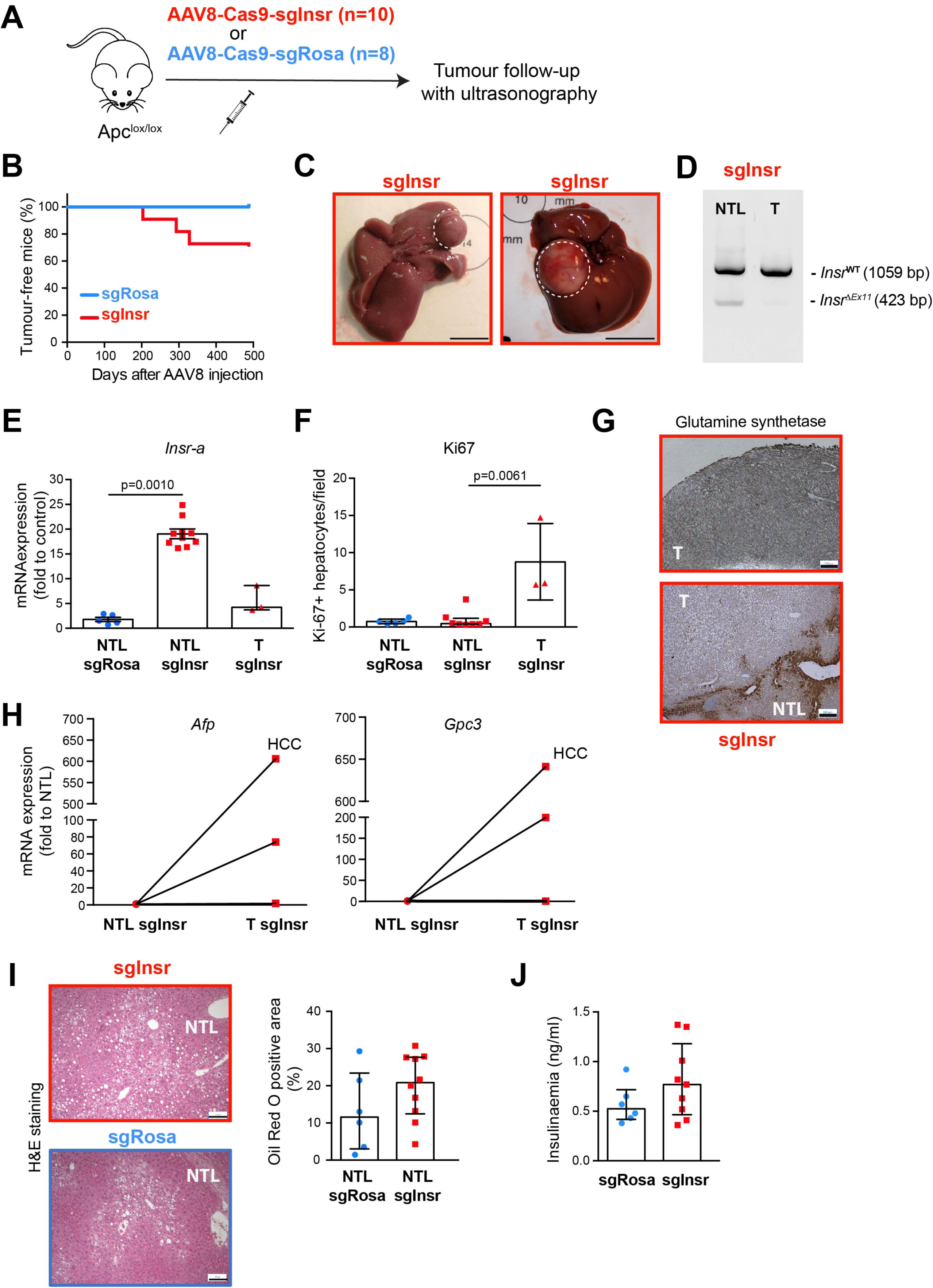
Expression of INSR-A in hepatocytes leads to tumour development in the long term. Mice were injected with recombinant AAV8 encoding saCas9 and sgRNA against *Rosa* (sgRosa, n=8) or *Insr* (sgInsr, n=10) locus and followed by ultrasonography during 13 months before sacrifice. **A.** Schematic illustration of the protocol. **B.** Tumour appearance over time. **C.** Representative images of two sgInsr livers at sacrifice. Tumours are surrounded by a dotted line. Scale bar: 1 cm. **D.** Representative image of *Insr* gene edition evaluated by PCR in nontumour liver (NTL) and tumour (T) tissues. **E.** Expression of *Insr-a* transcripts in nontumour livers (NTL) and tumours (T) evaluated by RT-qPCR (n=3-10). **F.** Quantification of Ki67-positive proliferating hepatocytes by immunohistochemistry (n=3-10). **G.** Representative images of glutamine synthetase staining in two sgInsr tumours (T) and adjacent non-tumour liver (NTL). **H.** Expression of *Afp* and *Gpc3* transcripts in three sgInsr tumours (T) and their adjacent nontumour liver (NTL) evaluated by RT-qPCR. **I.** Representative images of H&E staining of non-tumour sgRosa and sgInsr livers (NTL). **J.** Quantification of steatosis by oil red O staining (n=6-10). **K.** Insulinaemia measured on 5-6 h fasted mice (n=6-9). Data are median ± IQR.

### Expression of INSR-A in hepatocytes boosts β-catenin-driven liver carcinogenesis

To investigate the impact of INSR-A on oncogene-induced liver carcinogenesis, we first used Apc^lox/lox^ mice in which the injection of adenovirus encoding Cre recombinase (AdCre) results in the specific loss of *Apc* in a fraction of hepatocytes (APC^Δhep^ model) and the subsequent constitutive activation of β-catenin, leading to the emergence of two liver tumour types: (i) differentiated HCCs mimicking the human HCC subclasses G5 and G6; and (ii) undifferentiated tumours clustering with pediatric mesenchymal hepatoblastoma (HB).^27,28,36,37^ In APC^Δhep^ model, *Insr-a* mRNA and *Insr-a*:*Insr-b* mRNA ratio were substantially induced in HB-like tumours but not in HCCs as compared to NTL (**Figure S6A**). The total expression of *Insr* mRNA tended to be decreased in HB-like tumours (**Figure S6A**). To determine whether the mouse data were relevant to human HCC and HB, we analyzed RNA-seq datasets from large patient cohorts. We observed that : (i) the proportion of *INSR-A* mRNA was significantly higher in human HBs than in NTL, irrespective of subclass; (ii) the proportion of *INSR-A* mRNA was higher in human HCCs than in NTL, but to a lesser extent for HCC subclasses G5-G6; and (iii) *Insr* transcript was significantly decreased in all three HB subclasses compared with NTL, while its expression was slightly decreased in HCC subclass G6, but not in subclass G5 (**Figure S6B**).

Apc^lox/lox^ mice were injected with AAV8-sgInsr (n=8) or AAV8-sgRosa (n=6), then with AdCre and were subsequently monitored for tumour development (**Figure 3A**). While the APC^Δhep^ mice injected with AAV8-sgRosa did not develop tumours until day 125, it took only 45 days for the first APC^Δhep^ mouse injected with AAV8-sgInsr to develop tumours (**Figure 3B**). Ultrasound measurements of the liver performed in these mice as well as in an additional cohort (5 sgRosa/APC^Δhep^ and 5 sgInsr/APC^Δhep^ mice) revealed that cumulative tumour volumes were significantly higher at day 200 in sgInsr/APC^Δhep^ mice than in sgRosa/APC^Δhep^ mice (**Figure 3C**). However, comparison of the growth rate of individual tumours assessed from ultrasound measurements showed that there was no difference in the progression of sgRosa/APC^Δhep^ and sgInsr/APC^Δhep^ tumours (**Figure S7A**). At the time of sacrifice, 46% of sgInsr/APC^Δhep^ mice (6 out of 13) had enlarged livers with macroscopic nodules too numerous to be counted (**Figure 3D**). As expected, NTL from sgInsr/APC^Δhep^ mice showed an increased expression of *Insr-a* mRNA compared to NTL from sgRosa/APC^Δhep^ mice (**Figure 3E**). *Apc* mRNA excision was similar in sgRosa/APC^Δhep^ and sgInsr/APC^Δhep^ tumours (**Figure 3F**), indicating that the β-catenin pathway was activated in a comparable manner in both groups. However, we observed that the presence of INSR-A in the liver parenchyma significantly altered the proportion of HCC and HB-like tumours as assessed by histology and immunohistochemical staining for GS in macro- and micro-nodules. Thus, a higher percentage of differentiated HCC (85%) was detected in sgInsr/APC^Δhep^ livers than in sgRosa/APC^Δhep^ livers (44%) (p<0.0001) (**Figure 3G**). Tumour transcriptome analysis revealed no significant difference between the sgRosa/APC^Δhep^ and sgInsr/APC^Δhep^ groups for each tumour type (**Figure 3H**). HB-like tumours but not HCC had high levels of *Insr-a* transcript, irrespective of the liver from which they arose (**Figure S7B**). Finally, *Insr* gene editing was analyzed and showed that, unlike sgInsr/APC^Δhep^ NTL, the vast majority of sgInsr/APC^Δhep^ tumours (93.5%; 29/31) was not edited at the *Insr* locus (**Figure 3I**), mirroring what had been observed with spontaneous tumours (**Figure 2D**). This indicates that the *Insr-a* mRNA detected in some sgInsr/APC^Δhep^ tumours, particularly in HB-like tumours, was not the result of *Insr* gene editing but of a secondary alteration in alternative mRNA splicing during tumour progression.

**Figure 3:**
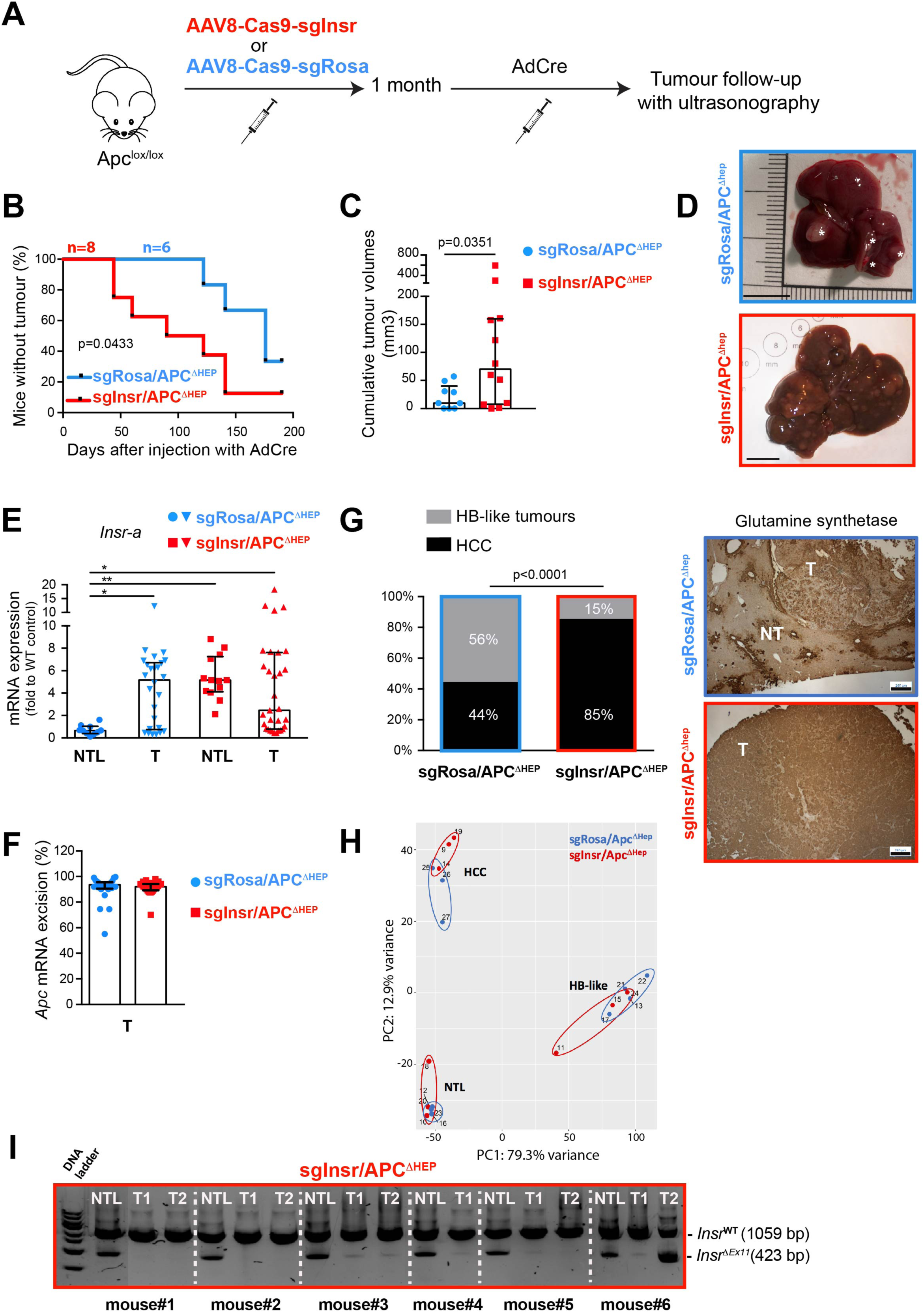
Expression of INSR-A in hepatocytes boosts β-catenin-driven liver carcinogenesis. Apc^lox/lox^ mice were injected with recombinant AAV8 encoding saCas9 and sgRNA against *Rosa26* (sgRosa/APC^ΔHep^, n=6) or *Insr* (sgInsr/APC^ΔHep^, n=8) locus. One month later, mice were injected with an adenovirus encoding the Cre recombinase (AdenoCre) and followed by ultrasonography. Mice were sacrificed before cumulative tumour diameter reached 1.2 cm. **A.** Schematic illustration of the protocol. **B.** Tumour appearance over time. **C.** Cumulative tumour volumes evaluated by ultrasonography 200 days after AdenoCre injection (n=9-12). Note that one sgInsr/APC^ΔHep^ liver could not be evaluated because too many tumors were detected at the time of ultrasound. **D.** Macroscopic pictures of representative livers from sgRosa/APC^ΔHep^ and sgInsr/APC^ΔHep^ mice at sacrifice. Scale bar: 1 cm. **E.** Expression of *Insr-a* mRNA in non-tumour livers (NTL) and tumours (T) evaluated by RT-qPCR (n=10-28). **F.** Expression of excised *Apc* mRNA in tumours (T) evaluated by RT-qPCR (n=25-28). **G.** Percent of microscopically and macroscopically detected HCC and HB-like tumours in sgRosa/APC^ΔHep^ and sgRosa/APC^ΔHep^ livers (*left panel*). Representative images of glutamine synthetase staining in sgRosa/APC^ΔHep^ HB and sgInsr/APC^ΔHep^ HCC. Scale bar: 260 μM (*right panel*)**. H.** Principal component analysis of the transcriptome from sgRosa/APC^ΔHep^ and sgInsr/APC^ΔHep^ non-tumour livers (NTL) and tumours (n=3 for each group). **I.** Representative picture of *Insr* gene edition evaluated by PCR in nontumour livers (NTL) and tumours (T) from sgInsr/APC^ΔHep^ mice. Only tumour T2 in mouse#6 was edited at *Insr* locus. Data are median ± IQR.

To strengthen these data, we used a second mouse model of β-catenin-mediated liver cancer in which sustained β-catenin activation was induced in hepatocytes by CRISPR/Cas9-mediated loss of *Ctnnb1* exon 3 (βCAT^Δex3^ model).^28^ AAV8 encoding sgRNA against *Ctnnb1* were injected together with those targeting *Insr* or *Rosa* locus and tumour development was monitored by ultrasonography (**Figure S7C**). We observed an earlier occurrence of tumour initiation in sgInsr/βCAT^Δex3^ mice (**Figure S7D**), along with increased cumulative tumour volumes at day 300 compared with sgRosa/βCAT^Δex3^ mice (**Figure S7E**). NTL from sgInsr/βCAT^Δex3^ mice overexpressed *Insr-a* transcript (**Figure S7F**) and were edited at the *Insr* locus (**Figure S7G**). Tumours in each group were mainly differentiated HCC, with the exception of one sgRosa/βCAT^Δex3^ tumour which was undifferentiated (data not shown). sgRosa/βCAT^Δex3^ and sgInsr/βCAT^Δex3^ tumours had similar levels of deleted *Ctnnb1* mRNA (**Figure S7H**). As in the case of spontaneous sgInsr tumours and sgInsr/APC^Δhep^ tumours, the vast majority of sgInsr/βCAT^Δex3^ tumours (72.7%; 8/11) were not edited at the *Insr* locus (**Figure S7G**).

In both genetic models of β-catenin-driven liver carcinogenesis, there was no difference in fasting glycaemia and insulinaemia at sacrifice between mice with hepatic expression of INSR-A and control mice (**Figures S7I** and **S7J**). Taken together, our data indicate that the presence of hepatocytes expressing INSR-A accelerates β-catenin-driven liver carcinogenesis, even though they do not themselves engage in this process. This is true whether *Insr* gene editing is performed before (APC^Δhep^ model) or at the same time (βCAT^Δex3^ model) as activation of the β-catenin pathway. The promotion of liver cancer initiated by INSR-A occurred in the absence of major disturbances of glucose homeostasis. Moreover, the presence of INSR-A in liver parenchyma also seems to be a determinant for the commitment of β-catenin-activated hepatocyte towards HCC rather than HB.

### Expression of INSR-A in the liver parenchyma from APC^Δhep^ mice generates inflammation, proliferation and apoptosis

To investigate the mechanisms underlying the acceleration of β-catenin driven hepatocarcinogenesis by hepatic expression of INSR-A, sgRosa/APC^Δhep^ and sgInsr/APC^Δhep^ mice were sacrificed 50 days after AdCre injection, at an early step preceding macroscopic tumour formation (**Figure 4A**). At sacrifice, two out of five sgInsr/APC^Δhep^ mice showed significant hepatomegaly (**Figure 4B**) together with elevated levels of the liver-specific enzymes AST and ALT, which are indicative of liver damage (**Figure S8A**). Histopathological examination of H&E sections revealed the presence of microvacuolar steatosis and glycogenated nuclei in all sgInsr/APC^Δhep^ livers (**Figure 4C**). Necroinflammation with apoptotic bodies was also observed in sgInsr/APC^Δhep^ livers along with the presence of inflammatory cells disseminated in the whole liver. Accordingly, myeloperoxidase (MPO) staining, which is a marker for neutrophil and monocyte granules, was significantly increased in sgInsr/APC^Δhep^ livers (**Figure 4D**). Immunohistochemical staining of F4/80, a marker of macrophages, was also increased in sgInsr/APC^Δhep^ livers (**Figure 4E**). Consistent with the above observations, the expression of cell-surface myeloid markers Ly6C (for bone marrow-derived monocytes and neutrophils) and Ly6G (for mature neutrophils) was induced in sgInsr/APC^Δhep^ livers compared with sgRosa/APC^Δhep^ livers (**Figure S8B**). To further characterize inflammatory mediators, a proteome profiler cytokine/chemokine array was performed on whole-liver protein lysates (**Figure 4F**). Several signals were upregulated in protein extracts from sgInsr/APC^Δhep^ livers compared with those from sgRosa/APC^Δhep^ livers, and those signals which showed the highest levels of induction were related to myeloid cell accumulation: MPO, lipocalin 2 (LCN2, also named neutrophil gelatinase associated lipocalin (NGAL), which induces neutrophil attracting signals in the injured liver ^38^), MMP9 (a gelatinase which is released primarily by neutrophils and monocytes, and has chemotactic effect on neutrophils possibly *via* its binding to LCN2 ^39^), chitinase 3-like 1 (CHI3L1, which is highly expressed in activated neutrophils and macrophages and promote the secretion of chemoattractants ^40–42^), osteopontin (OPN, encoded by *Spp1*, secreted by macrophages, and involved in neutrophil attraction in relation with apoptosis ^43^), and LDL receptor (recently shown to be involved in neutrophil-to-hepatocyte communication ^44^). Gene expression analysis of *Lcn2*, *Chi3l1*, and *Spp1* confirmed these data (**Figure S8C**). Higher expression of *Ccl2, Csf1*, and *Cxcl2* mRNA (encoding chemokines that orchestrate myeloid cells interaction and recruitment during liver inflammation ^45^), as well as higher expression of the pro-inflammatory cytokine TNF-α were observed in sgInsr/APC^Δhep^ livers (**Figure 4G**). These data led us to question the impact of this inflammatory environment on cell death and proliferation. Apoptosis was studied using the TUNEL method and IHC detection of cleaved caspase-3 in hepatocytes. Both techniques revealed a significant increase in cell death in the livers of sgInsr/APC^Δhep^ mice, compared with sgRosa/APC^Δhep^ livers (**Figures 4H** and **S8D**). In parallel, increased hepatocyte proliferation was observed in sgInsr/Apc^Δhep^ livers compared with sgRosa/APC^Δhep^ livers, as assessed by Ki67 IHC staining and RT-QPCR (**Figure S8E**).

**Figure 4:**
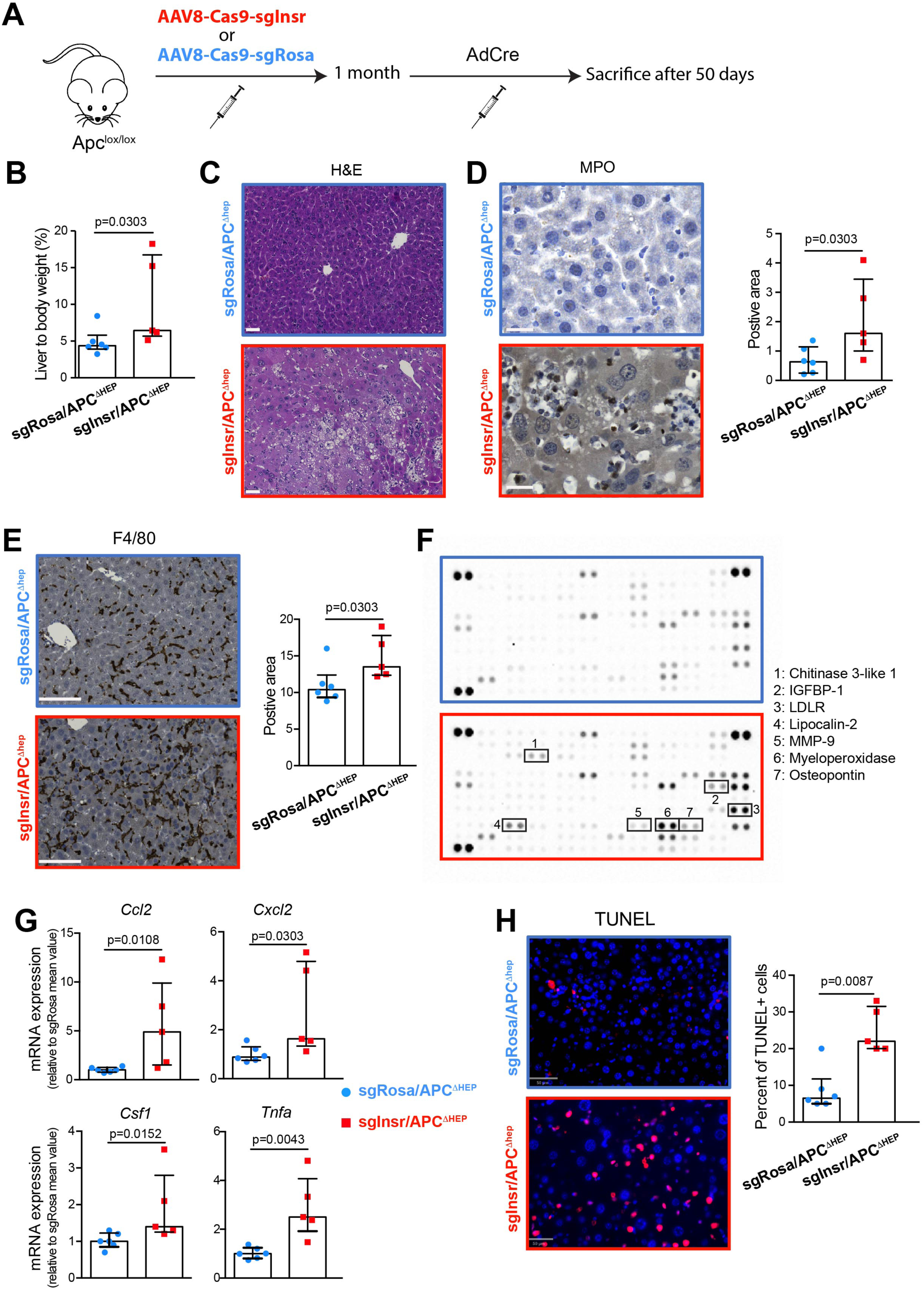
Expression of INSR-A generates inflammation and apoptosis in APC^ΔHep^ livers at day 50. Apc^lox/lox^ mice were injected with recombinant AAV8 encoding saCas9 and sgRNA against *Rosa26* (sgRosa/APC^ΔHep^, n=6) or *Insr* (sgInsr/APC^ΔHep^, n=5) locus. One month later, mice were injected with an adenovirus encoding the Cre recombinase (AdenoCre) and sacrificed 50 days later. **A.** Schematic illustration of the protocol. **B.** Liver-to-body weight ratio (n=5-6). **C.** Representative images of H&E staining showing microvacuolar steatosis, glycogenated nuclei, apoptotic bodies and inflammatory cells in sgInsr/APC^Δhep^ livers. Scale bar: 50 μM. **D.** Representative images of myeloperoxidase (MPO) IHC staining from sgRosa/APC^Δhep^ and sgInsr/APC^Δhep^ livers (*left panel*, scale bar: 120 μM); Quantification of MPO staining (*right panel*; n=5-6). **E.** Representative images of F4/80 IHC staining from sgRosa/APC^Δhep^ and sgInsr/APC^Δhep^ livers (*left panel*, scale bar: 100 μM); Quantification of F4/80-positive macrophages (*right panel*; n=5-6). **F.** Representative images of proteome profiler cytokine/chemokine arrays performed on whole-cell lysates from sgRosa/APC^Δhep^ and sgInsr/Apc^Δhep^ livers (n=2). **G.** Expression of transcripts encoding chemokines/cytokines evaluated by RT-qPCR (n=5-6). **H.** Representative images of TUNEL assay from sgRosa/APC^Δhep^ and sgInsr/APC^Δhep^ livers (*left panel*); Quantification of TUNEL-positive cells (*right panel*, n=5-6). Data are median ± IQR.

### The presence of Insr-deleted hepatocytes stimulates the expansion of APC^Δhep^ hepatocytes while disappearing from liver parenchyma

Interestingly, most of the cytokine and chemokine markers overexpressed in sgInsr/APC^Δhep^ livers, had been identified as being induced at transcriptional level in livers fully activated for the β-catenin pathway (from Apc^lox/lox^-CreER^T2^ mice injected with a high dose tamoxifen) compared with WT livers **(Figure S8F)**, suggesting that APC^Δhep^-dependent gene expression program could be favoured in sgInsr/APC^Δhep^ livers. Therefore, the status of hepatocytes genetically modified to activate the β-catenin pathway was investigated in sgInsr/APC^Δhep^ livers. The expression of *Glul*/GS and Axin 2, two direct β-catenin targets, evaluated by RT-QPCR and/or IHC, was significantly induced in sgInsr/APC^Δhep^ livers, by comparison with sgRosa/APC^Δhep^ livers (**Figures 5A** and **5B**). This can be taken as the result of APC^Δhep^ hepatocyte proliferation, given that the percentage of excised *Apc* mRNA was significantly higher in sgInsr/APC^Δhep^ livers (**Figure 5C**) and that there was a strong positive correlation between *Glul* and excised *Apc* mRNA levels (**Figure 5D**). Moreover, the two livers with the highest hepatomegaly (**Figure 4B**) were those with the highest *Apc* mRNA excision. It is noteworthy that fasting blood glucose was significantly reduced in sgInsr/APC^Δhep^ mice at day 50 (**Figure 5E**), which may be consistent with the repressive action of β-catenin on gluconeogenesis in the liver.^46^ To characterize the gene signatures expressed by APC^Δhep^ hepatocytes from sgInsr/APC^Δhep^ livers, sgRosa/Apc^lox/lox^ and sgInsr/Apc^lox/lox^ mice were given a high dose of AdCre-GFP injection, and GFP-positive hepatocytes were sorted by flow cytometry on day 21, as previously described.^47^ Transcriptomic analysis performed on the sorted cells revealed no differences between APC^KO^ hepatocytes isolated from sgRosa/APC^Δhep^ and those isolated from sgInsr/APC^Δhep^ livers (data not shown). These data suggest that forced expression of INSR-A does not affect the intrinsic transcriptional program initiated by oncogenic β-catenin in APC^Δhep^ hepatocytes although its does promote their expansion.

**Figure 5:**
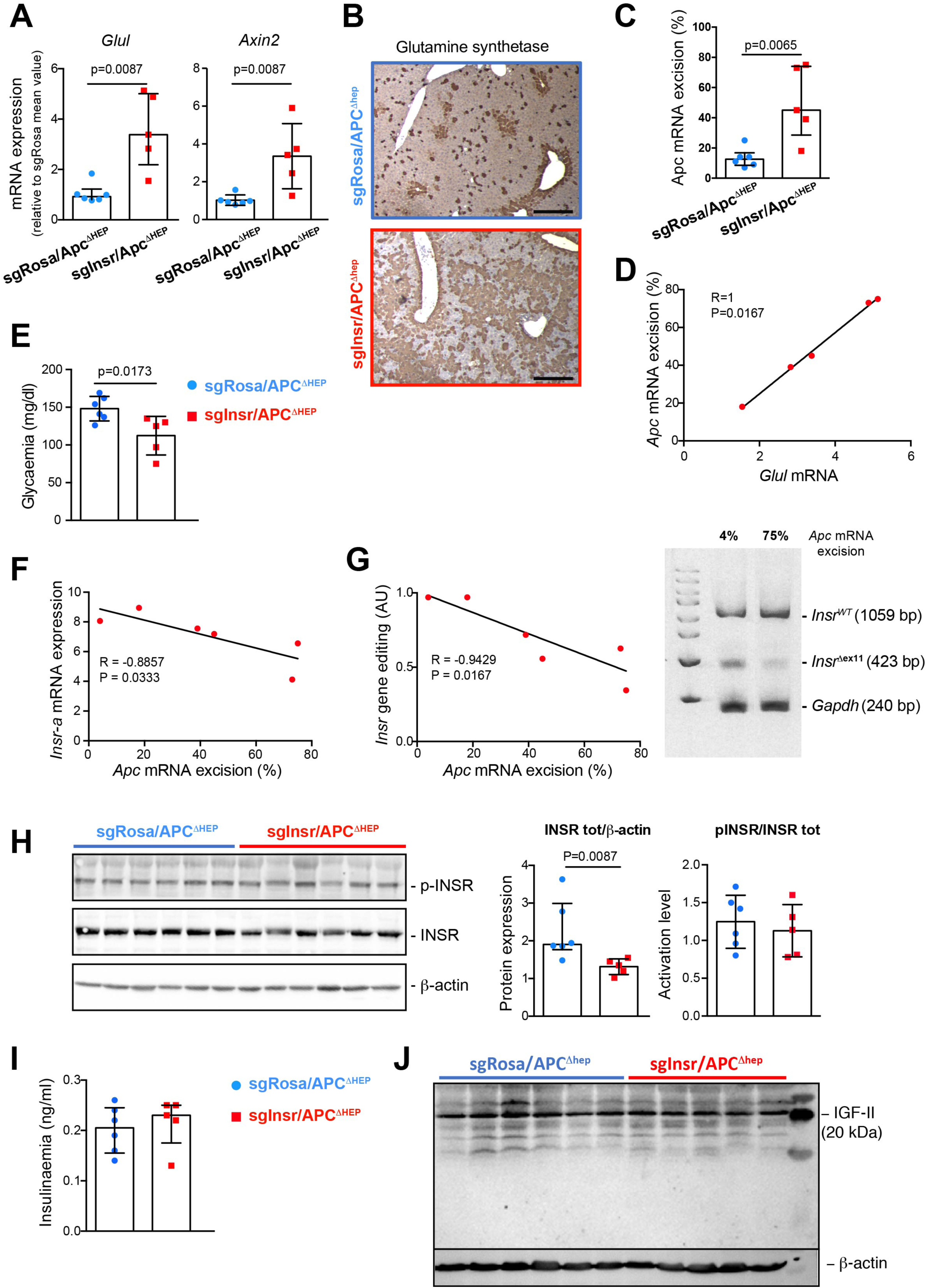
INSR-A expression is not maintained in APC^ΔHep^ livers at day 50. Apc^lox/lox^ mice were injected with recombinant AAV8 encoding saCas9 and sgRNA against *Rosa26* (sgRosa/APC^ΔHep^, n=6) or *Insr* (sgInsr/APC^ΔHep^, n=5) locus. One month later, mice were injected with an adenovirus encoding the Cre recombinase (AdenoCre) and sacrificed 50 days later. **A.** Expression of *Glul* and *Axin2* transcripts evaluated by RT-qPCR (n=5-6). **B.** Representative images of glutamine synthetase IHC staining from sgRosa/APC^Δhep^ and sgInsr/APC^Δhep^ livers; scale bar: 260 μM. **C.** Expression of excised *Apc* mRNA evaluated by RT-qPCR (n=5-6). **D.** Correlation between excised *Apc* and *Glul* mRNA. Spearman’s correlation coefficient is indicated. **E.** Fasting glycaemia measured on 5-6 h fasted mice (n=5-6). **F.** Correlation between excised *Apc* and *Insr-a* mRNA. Spearman’s correlation coefficient is indicated. **G.** Correlation between excised *Apc* mRNA and *Insr* gene edition. Spearman’s correlation coefficient is indicated (*left panel*). A representative image of *Insr* gene editing evaluation is shown (*right panel*). Numbers above the gel indicate the percentage of *Apc* mRNA excision. **H.** Western blot analysis of phospho/total INSR protein levels on whole-cell lysates from sgRosa/APC^ΔHep^ (n=6) and sgInsr/APC^ΔHep^ (n=5) livers. β-actin detection is used as a loading control (*left panel*). Quantification by scanning densitometry (*right panel*). **I.** Plasma insulinaemia measured on 5-6-h fasted mice (n=5-6). **J.** Western blot analysis of IGF-II protein levels on whole-cell lysates from sgRosa/APC^ΔHep^ (n=6) and sgInsr/APC^ΔHep^ (n=5) livers. β-actin detection is used as a loading control. Data are median ± IQR.

The fact that the hepatocytes which had been genetically edited to express INSR-A did not engage into the transformation pathway then led us to examine their fate in sgInsr/APC^Δhep^ livers. As shown in **Figure 5F**, there was an inverse correlation between the levels of *Apc* mRNA excision and *Insr-a* mRNA in sgInsr/APC^Δhep^ livers. Since *Insr-a* mRNA could originate not only from genetically-modified hepatocytes, but also from other liver cell types, such as immune cells that express it endogenously ^48^, we also evaluated the presence of edited hepatocytes in sgInsr/APC^Δhep^ liver parenchyma by detecting genomic DNA deletion with semi-quantitative PCR. Consistent with the analysis conducted at the mRNA level, a negative correlation was observed between *Apc* mRNA excision and DNA edition at the *Insr* locus (**Figure 5G**), indicating that the more the β-catenin pathway was activated in the liver, the fewer hepatocytes were edited in the *Insr* locus. Furthermore, INSR protein content was significantly reduced in sgInsr/APC^Δhep^ livers compared with sgRosa/APC^Δhep^ livers. Levels of activated INSR and its ligands, insulin and IGF-II, were comparable between sgRosa/APC^Δhep^ and sgInsr/APC^Δhep^ livers (**Figures 5H-5I**). Taken together, these data suggest that the promotion of oncogenic β-catenin signalling in sgInsr/APC^Δhep^ liver parenchyma is accompanied by downregulation of the INSR pathway, as evidenced by the disappearance of *Insr*-edited hepatocytes and reduced INSR content, in a context of apoptosis.

### INSR-A expressing hepatocytes are more sensitive to apoptosis in vitro

Prominent cell death, disappearance of edited hepatocytes and INSR down regulation in sgInsr/APC^Δhep^ livers led us to assess the intrinsic susceptibility to apoptosis of murine hepatocytes expressing INSR-A. To this end, hepatocytes were isolated from sgRosa and sgInsr mice by collagenase liver perfusion and cultured in 2D. *In vitro* culture maintained INSR-A signaling for at least 24 h, as sgInsr hepatocytes overexpressed *Insr-a* mRNA (**Figure 6A**) and were more sensitive to IGF-II for INSR and AKT activation, compared with sgRosa hepatocytes, while the cellular response to insulin was similar in both groups (**Figure 6B**). A high dose of staurosporine and a low dose of staurosporine combined with TNF-α were used as pro-apoptotic inducers as previously reported.^49^ Hepatocytes were deprived of growth factors overnight and washed before being treated for 24 h with the pro-apoptotic drugs. Both treatments induced significantly greater apoptosis in sgInsr hepatocytes than in sgRosa hepatocytes, as assessed by induction of PARP and caspase-3 cleavages (**Figure 6C**). Hepatocyte death resulted concomitantly in the downregulation of INSR protein expression. However, no differential effect was observed between sgRosa and sgInsr hepatocytes (**Figure 6C**). This *in vitro* approach demonstrates that INSR-A possesses an increased intrinsic capacity to relay pro-apoptotic signals in hepatocytes.

**Figure 6:**
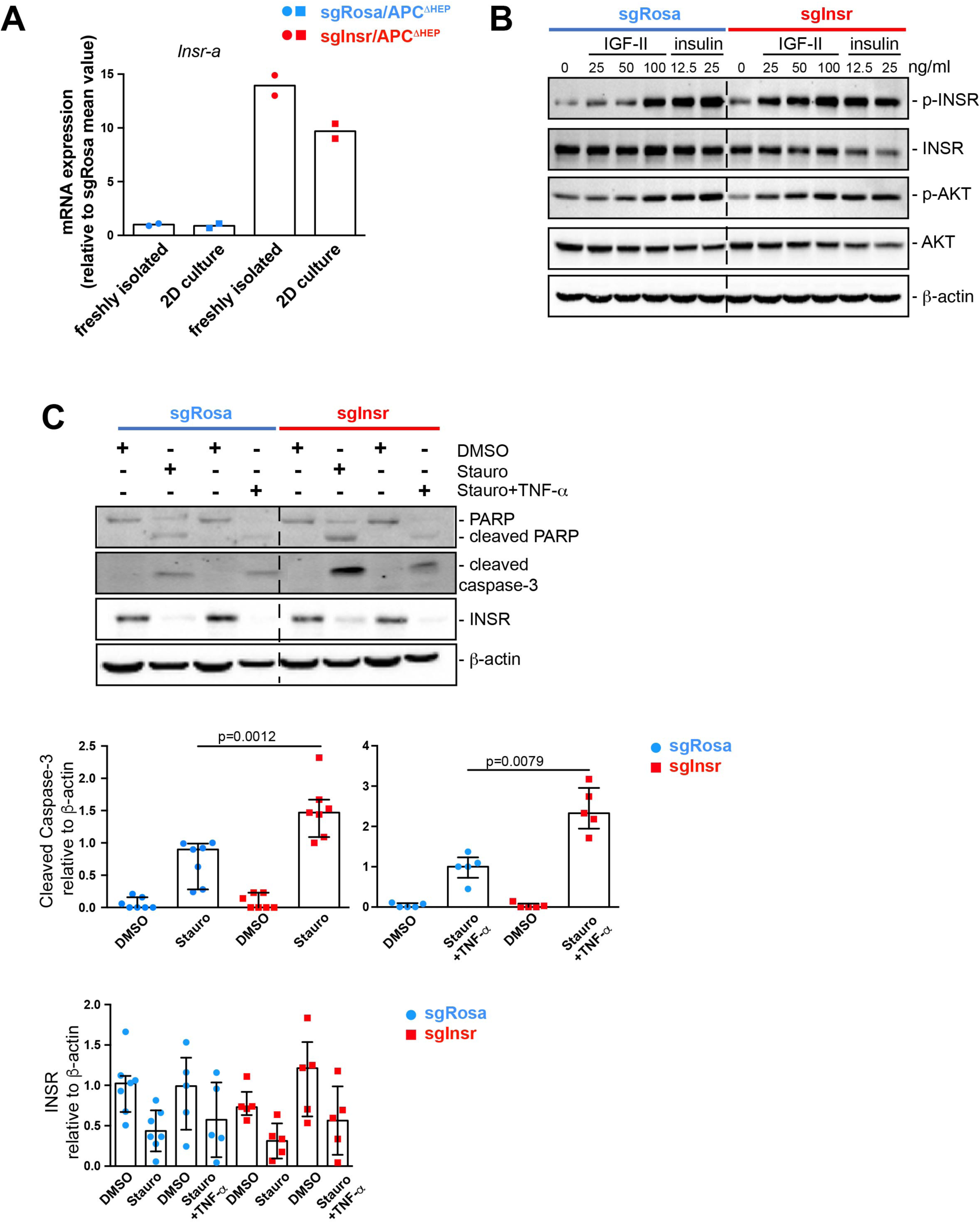
INSR-A expressing hepatocytes exhibit intrinsic high sensitivity to pro-apoptotic signals *in vitro*. Mice were injected with recombinant AAV8 encoding saCas9 and sgRNA against *Rosa26* (sgRosa) or *Insr* (sgInsr) and sacrificed one month later. **A.** Expression of *Insr*-a transcript by RT-qPCR in freshly isolated hepatocytes and after 24 h in 2D culture. **B.** Representative image of Western blot analysis of phospho/total INSR and AKT protein levels on whole-cell lysates from sgRosa (n=4) and sgInsr (n=4) hepatocytes. Cells were stimulated with insulin or IGF-II during 10 min. β-actin detection is used as a loading control. **C.** Representative image of Western blot analysis of PARP, cleaved caspase-3 and INSR protein levels on whole-cell lysates from sgRosa (n=5-7) and sgInsr (n=5-7) hepatocytes. Cells were stimulated with staurosporine (Stauro, 10 μM) or Staurosporine (3.33 μM) plus TNF-α (7 ng/ml) during 24 h. β-actin detection is used as a loading control (*Upper panel*). Quantification by scanning densitometry (*middle and lower panels*).

## DISCUSSION

A large body of *in vitro* data has helped characterize INSR-A properties in a variety of carcinoma cell lines. In HCC, the presence of high INSR-A expression has been associated with the acquisition of markers of aggressiveness, tumour progression, and poor prognosis in patients undergoing surgery.^13,14,17^, ^present^ ^study^ The identification that splicing machinery defects can occur very early in the course of chronic liver diseases ^17,50,51^, and as a consequence INSR-A is expressed in pathological livers, raises questions about the impact on liver homeostasis and tumour initiation of INSR-A. Using CRISPR/Cas9 genetically engineered mice, we directly explored the importance of INSR-A in liver homeostasis and hepatic tumour initiation by replacing INSR-B with INSR-A in a fraction of hepatocytes.

Although INSR-B and INSR-A have similar affinity for insulin *in vitro*, discrepancies have been reported concerning their endocytosis kinetics and the nature of the intracellular signals they initiate downstream.^10,21^ Here, we did not observe any disturbance of carbohydrate homeostasis in sgInsr mice as assessed with GTT and ITT, nor any abnormal response of sgInsr hepatocytes to insulin stimulation *in vivo* and *in vitro*. Several hypotheses can be put forward for this. Given that the CRISPR/Cas9 system we have developed does not target all hepatocytes, it is possible that the number of non-genetically modified hepatocytes expressing INSR-B may be sufficient to transmit a normal biological response to insulin. Furthermore, it cannot be ruled out that INSR-A may share metabolic functions with INSR-B. Indeed, in mouse immortalized LIRKO hepatocytes, the overexpression of INSR-A induced an increase in insulin signalling that was associated with elevated glycogen synthesis and storage.^52^ Finally, insulin control of hepatic glucose production is complex, and some studies have questioned the contribution of hepatic IR to this process in mice and humans.^3,53–55^ Therefore, the alteration of INSR isoform relative expression may not have a major impact on hepatic glucose production in sgInsr mice. Further metabolic explorations in these mice are required to determine if other metabolic pathways are affected differentially in sgRosa and sgInsr mice. However, the presence of INSR-A in the liver parenchyma seems to disrupt homeostasis over the long term, given that some tumors have emerged in the sgInsr livers. The penetrance of the tumor phenotype is low, and it is difficult to speculate about the underlying mechanisms. It is conceivable that the pathophysiological mechanisms of aging (genomic instability, telomere dysfunction, epigenetic alterations, compromise of autophagy, mitochondrial dysfunction, cellular senescence, altered intercellular communication, deregulated nutrient sensing…) are integrated differently by hepatocytes genetically edited to express INSRA.

Very unexpectedly, the vast majority of tumours, whether occurring spontaneously or following activation of the β-catenin pathway, were not edited in the *Insr* gene, thus suggesting the counter-selection of INSR-A-expressing hepatocytes in the transformation process, and that INSR-A expression is not a selective advantage. A body of experimental evidence highlights the fact that the mechanism by which INSR-A modifies the environment in APC^ΔHep^ livers involves the entry of edited hepatocytes into apoptosis. Firstly, the presence of INSR-A is associated with hepatic inflammation, as evidenced by the accumulation of innate immune cells and the expression of chemokines in sgInsr/APC ^ΔHep^ livers, as compared with sgRosa/APC ^ΔHep^ livers. It is known that macrophages and neutrophils are responsible for clearing apoptotic hepatocytes.^56^ Both immune cell populations also cooperate and orchestrate the resolution of inflammation and tissue repair.^57^ In the sgInsr/APC ^ΔHep^ model, it would appear that liver inflammation is followed by tissue resolution in the following weeks, since transcriptomic analysis of liver tissue adjacent to tumours did not reveal signatures of fibrosis or chronic inflammation. Secondly, apoptosis was detected in sgInsr/APC ^ΔHep^ liver parenchyma by TUNEL and caspase-3 cleavage. While it may be difficult when using the TUNEL method to be sure that the DNA damage detected originates from apoptotic hepatocytes rather than from apoptotic immune cells, *i.e* neutrophils during the resolution of inflammation, the analysis of cleaved caspase-3 by IHC enabled us to clearly objectify hepatocyte apoptosis. Thirdly, the more the β-catenin pathway is activated, the less hepatocytes edited at *Insr* locus are maintained in the liver. This raises the question of what event(s) related to β-catenin signalling initiate(s) their disappearance. We have previously shown that the oncogenic activation of β-catenin triggers a specific inflammatory program leading to the remodeling of the liver microenvironment.^58^ This could prove deleterious to INSR-A expressing hepatocytes and promote their death, thus initiating a vicious circle of inflammation. Also of relevance here is the fact that β-catenin-driven inflammation in the liver does not involve the canonical inflammatory mediators of liver inflammation such as TNF-α.^58^ The fact that higher levels of TNF-α expression were evidenced in sgInsr/APC^Δhep^ livers indicates that the presence of INSR-A in liver parenchyma has modified liver inflammation qualitatively.

In both murine models of oncogenic β-catenin activation in the liver, INSR-A expression increased tumour onset and burden. Longitudinal monitoring of individual tumours revealed that once tumours appeared, INSR-A had no impact on tumour progression, indicating the likelihood that INSR-A influences the tumour initiation phases. We obtain no evidence that INSR-A affects the early transcriptional program initiated by oncogenic β-catenin in hepatocytes. Thus, one attractive hypothesis of how β-catenin-driven liver carcinogenesis may be promoted by INSR-A is cell competition. Cell competition is a vital mechanism that results in the elimination of less fit cells by their more fit neighbors.^59^ It acts as a quality control mechanism of tissue homeostasis during both developmental and adult stages. In addition, cell competition is used to remove early malignant cells from epithelial tissues. Cancer cells, however, can exploit it to promote tumour growth. In *Drosophila* midgut ^60^ and in murine intestinal organoids ^61–63^, APC^KO^ cells have been shown to acquire a crucial competitive advantage over surrounding wild-type cells, which are eliminated by different mechanisms thus allowing mutated cells to expand. In this context, it is tempting to consider that hepatocytes expressing INSR-A may be less adapted than hepatocytes expressing INSR-B to the signals emitted by APC^Δhep^ hepatocytes and will therefore be eliminated in their vicinity. The major expansion of β-catenin-activated hepatocytes detected only 50 days after AdCre injection in apoptotic sgInsr/Apc^Δhep^ livers supports this hypothesis.

The presence of INSR-A in the liver parenchyma not only boosts carcinogenesis in APC^Δhep^ mice but also reorients tumour fate by promoting the development of HCC rather than HB. Considering the increase in apoptosis in sgInsr/APC ^ΔHep^ livers, our data echo various studies that have shown that hepatocyte apoptosis generates a liver cytokine environment suitable to the development of HCC.^64,65^. These data will encourage us to investigate the molecular mechanisms by which inflammation linked to hepatocyte apoptosis affects the commitment of β-catenin activated hepatocytes in the transformation pathway leading to HCC.

*In vitro* culture of murine hepatocytes revealed that hepatocytes expressing INSR-A are intrinsically more susceptible to apoptosis than hepatocytes expressing INSR-B in the absence of ligand. Mechanistically, hepatocyte apoptosis was accompanied by downregulation of INSR protein expression, which may be due to increased receptor proteasomal degradation. Decreased INSR protein expression was observed consistently in apoptotic sgInsr/APC ^ΔHep^ livers. However, it was not possible to objectivate a significant difference in the decrease of INSR protein between apoptotic sgRosa and sgInsr hepatocytes *in vitro*, which could have accounted for their differential susceptibility to death. The molecular mechanisms underlying this observation have yet to be elucidated.

The identification of INSR-A as a receptor capable of transmitting a pro-apoptotic signal in the liver is suggestive of the TK activation-independent effects of INSR previously identified *in vitro* in different cell types.^66–69^ When unoccupied, INSR acts as a dependence receptor in mouse immortalized neonatal hepatocytes, brown preadipocytes and embryonic fibroblasts by increasing cell sensitivity to diverse pro-apoptotic signals. Interestingly, it has been reported that INSR-A overexpressed in mouse embryo fibroblasts is more effective than INSR-B in increasing the apoptotic cleavage of caspase-3 in response to DNA damage induced by etoposide.^68^ An expression imbalance between INSR-A and INSR-B could confer differential susceptibility to apoptosis in certain cells such as hepatocytes.

Taken as a whole, our work reveals a previously unrecognized function for INSR-A in liver tumour initiation related to its increased sensitivity to convey pro-apoptotic signals in pre-tumoral liver, a function that is radically different from that exerted during tumour progression. This work may be the first *in vivo* illustration of the involvement of a non-canonical ligand- and TK-independent function of an INSR isoform in pathology.

## Author contributions

C.D.-M. and F.L. conceived the study. C.D.-M. supervised all aspects of the project. F.L., A.C., C.G., S.S., A.A., R.A., and C.D.-M characterized sgInsr/APC^Δhep^, and sgInsr/βCAT^ΔEx3^. F.L, C.G. and A.I. performed histology, immunohistochemistry and Western blot analyses. F.L. and A.G. performed hepatocyte isolation and cell culture experiments. I.L. performed liver ultrasonography. S.P. validated CRISPR/Cas9 strategy *in vitro* and *in vivo*. J.A. performed liver histopathology analyses. S.I. and J.Z.R. provided human tumour data. A.G. managed Apc^loX/lox^ mouse breeding and provided samples. C.D.-M., A.G. and S.C. obtained funding supports. C.D.-M. wrote the manuscript with contributions from all authors.

## Acknowledgements

We are grateful to CRC core facilities (INSERM UMRS_1138, Sorbonne Université, Université Paris Cité) and their team members for animal care and breeding (CFE, Mrs Valérie Chauffeton and Mrs Sonia Prince), genotyping (CGB, Mrs Hermine Kakanakou), flow cytometry and cell imaging (CHICS, Dr Christophe Klein). We warmly thank the Institut Cochin’s core facilities (INSERM U1016, CNRS UMR8104, Université Paris Cité) and their team members for animal care and breeding (Moust’IC, Mr Marcio Do-Cruzeiro), liver ultrasonography (PIV, Dr Gilles Renault) and transcriptomic analysis (Genom’IC, Dr Franck Letourneur). The kidney function investigation facility at CRC is thanked for the plasma assays (Mrs Gaëlle Brideau). Dr Véronique Blouin (CAPACITES, INSERM UMR1089, Université de Nantes, France) is thanked for recombinant AAV8 production.

## Financial supports

This study has been supported by grants from INSERM, Ligue Nationale Contre le Cancer (to SC, team accreditation), Gefluc (to CDM), Fondation ARC pour la recherche sur le cancer (to CDM), and Institut National du Cancer (to AG, PLBIO 2021-095). Fanny Léandre was supported by grants from the Ministère de l’enseignement supérieur et de la Recherche and Ligue Nationale Contre le Cancer.

