## Supplemental data for "Hepatocyte expression of fetal insulin receptor isoform contributes to the promotion of liver cancer through non-cell autonomous mechanisms"

**Figure S1**

**A**

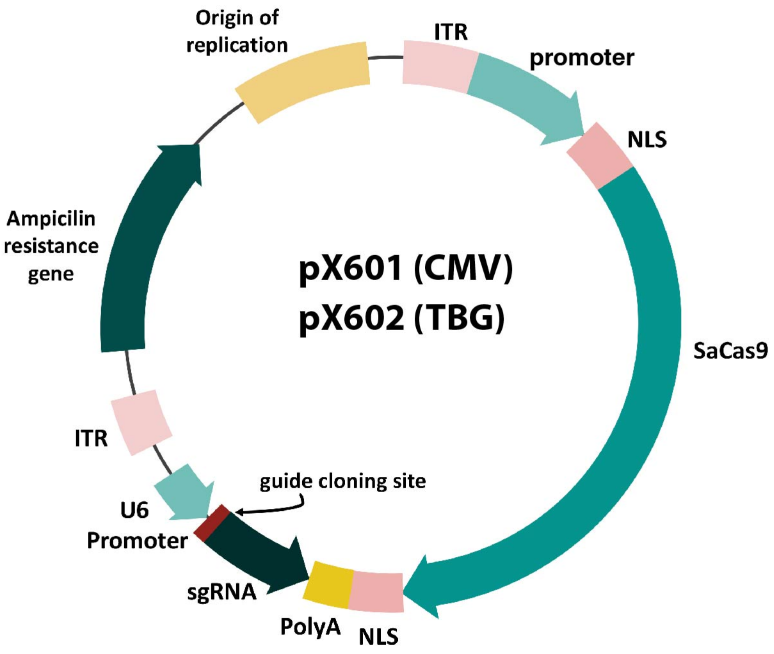
Origin of replication
ITR
promoter
NLS
Ampicilin resistance gene
pX601 (CMV)
pX602 (TBG)
SaCas9
ITR
guide cloning site
U6 Promoter
sgRNA
PolyA
NLS

**B**

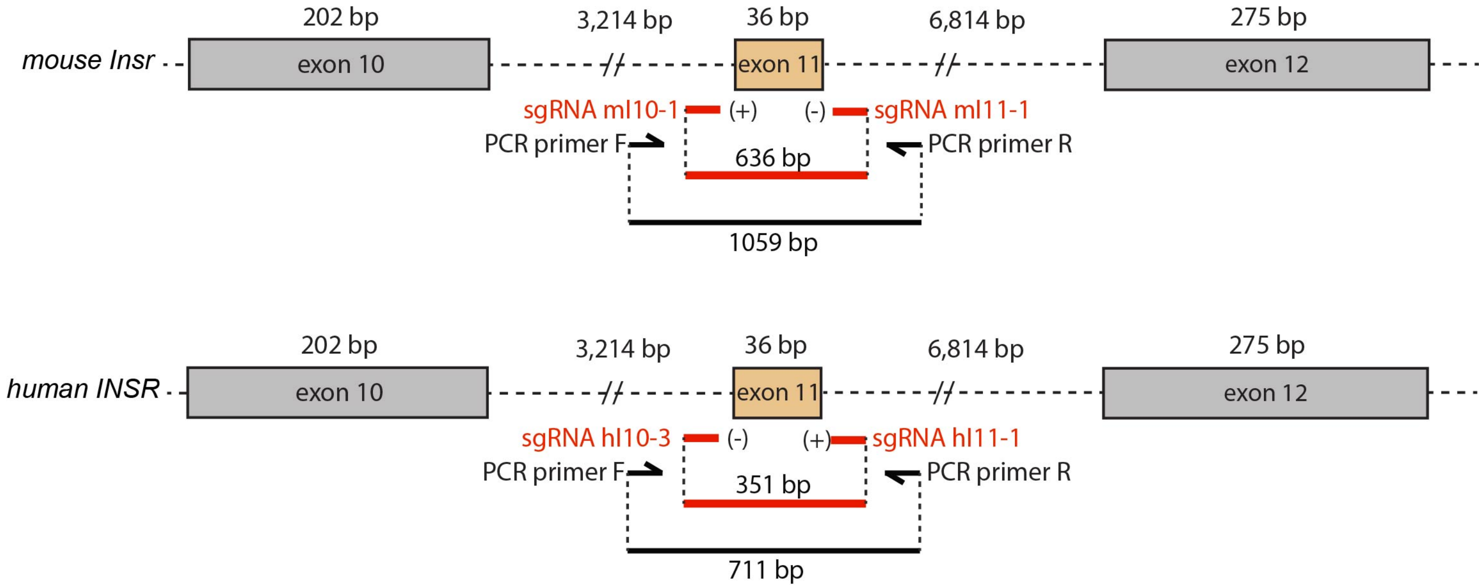
202 bp
3,214 bp
36 bp
6,814 bp
275 bp
mouse Insr
exon 10
exon 11
exon 12
sgRNA mI10-1
(+)
(-)
sgRNA mI11-1
PCR primer F
PCR primer R
636 bp
1059 bp
202 bp
3,214 bp
36 bp
6,814 bp
275 bp
human INSR
exon 10
exon 11
exon 12
sgRNA hI10-3
(-)
(+)
sgRNA hI11-1
PCR primer F
PCR primer R
351 bp
711 bp

### Figure S2

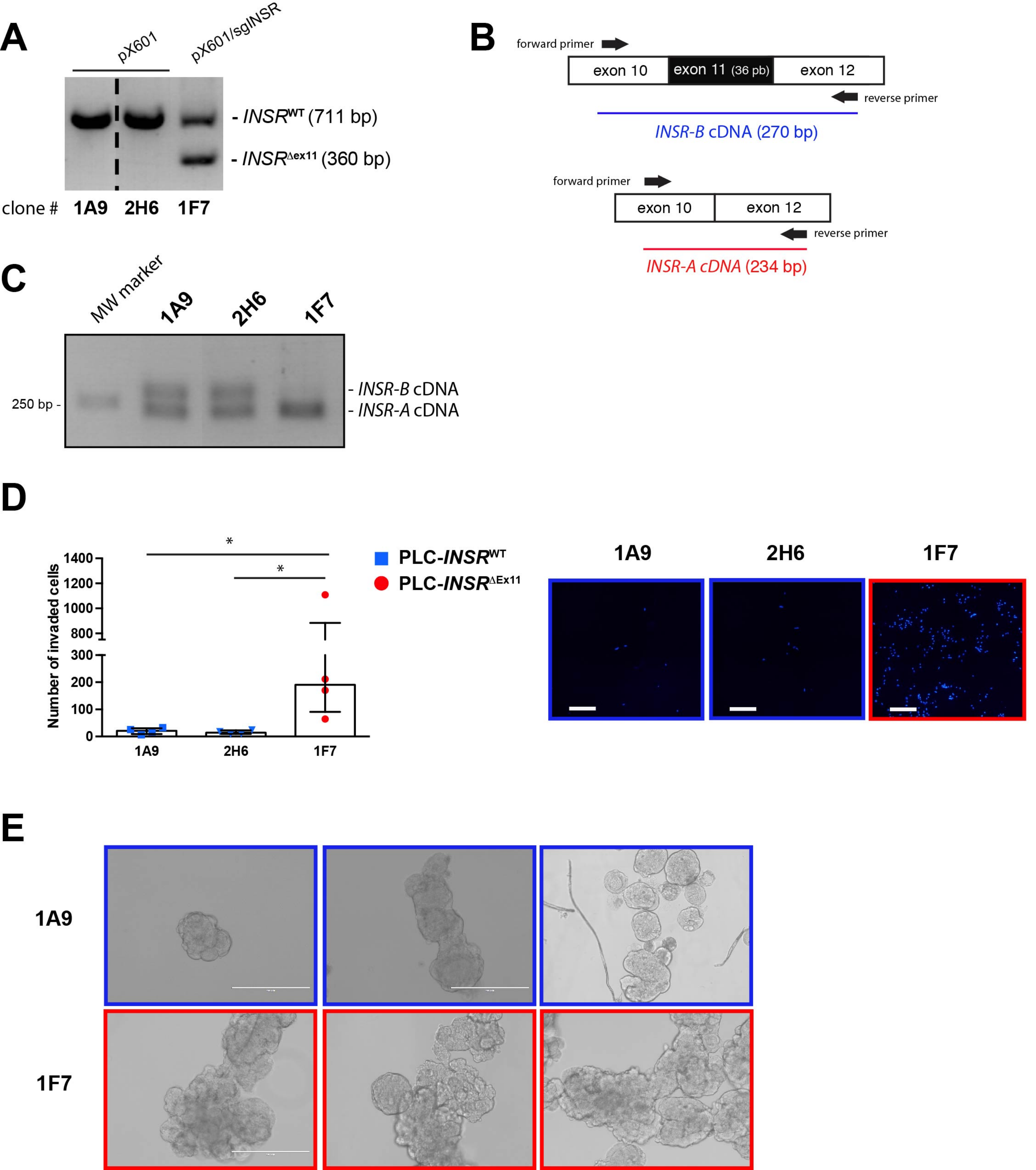
A
pX601
pX601/sgINSR
- INSRWT (711 bp)
- INSRΔex11 (360 bp)
clone # 1A9 2H6 1F7
B
forward primer
exon 10
exon 11 (36 pb)
exon 12
reverse primer
INSR-B cDNA (270 bp)
forward primer
exon 10
exon 12
reverse primer
INSR-A cDNA (234 bp)
C
MW marker
1A9
2H6
1F7
250 bp -
- INSR-B cDNA
- INSR-A cDNA
D
Number of invaded cells
1400
1200
1000
800
300
200
100
0
*
*
1A9
2H6
1F7
PLC-INSRWT
PLC-INSRΔEx11
1A9
2H6
1F7
E
1A9
1F7

### Figure S3

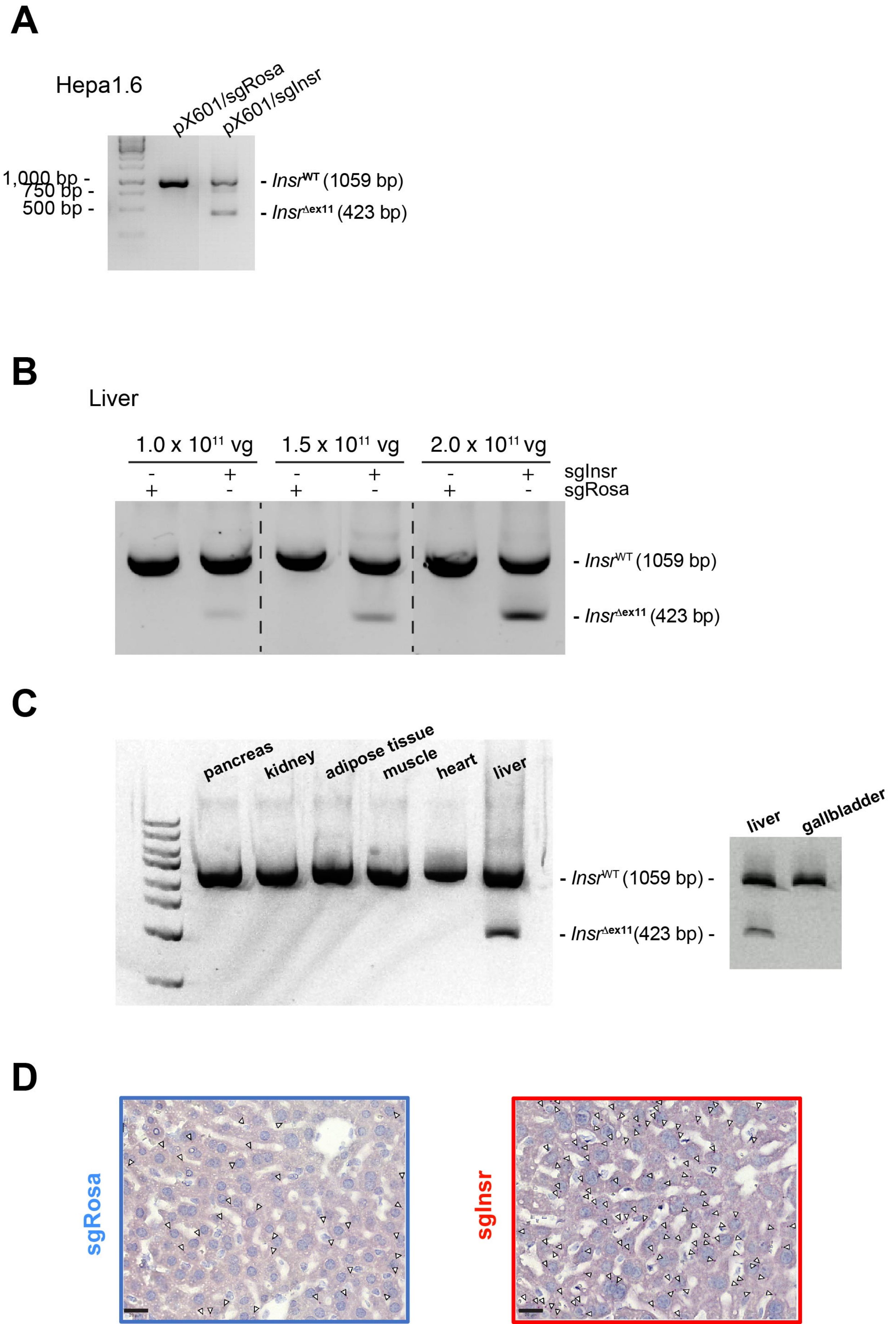

A
Hepa1.6
pX601/sgRosa
pX601/sgInsr
1,000 bp -
750 bp -
500 bp -
- Insr^WT (1059 bp)
- Insr^Δex11 (423 bp)
B
Liver
1.0 x 10^11 vg
1.5 x 10^11 vg
2.0 x 10^11 vg
sgInsr
sgRosa
- Insr^WT (1059 bp)
- Insr^Δex11 (423 bp)
C
pancreas
kidney
adipose tissue
muscle
heart
liver
- Insr^WT (1059 bp) -
- Insr^Δex11 (423 bp) -
liver
gallbladder
D
sgRosa
sgInsr

### Figure S4

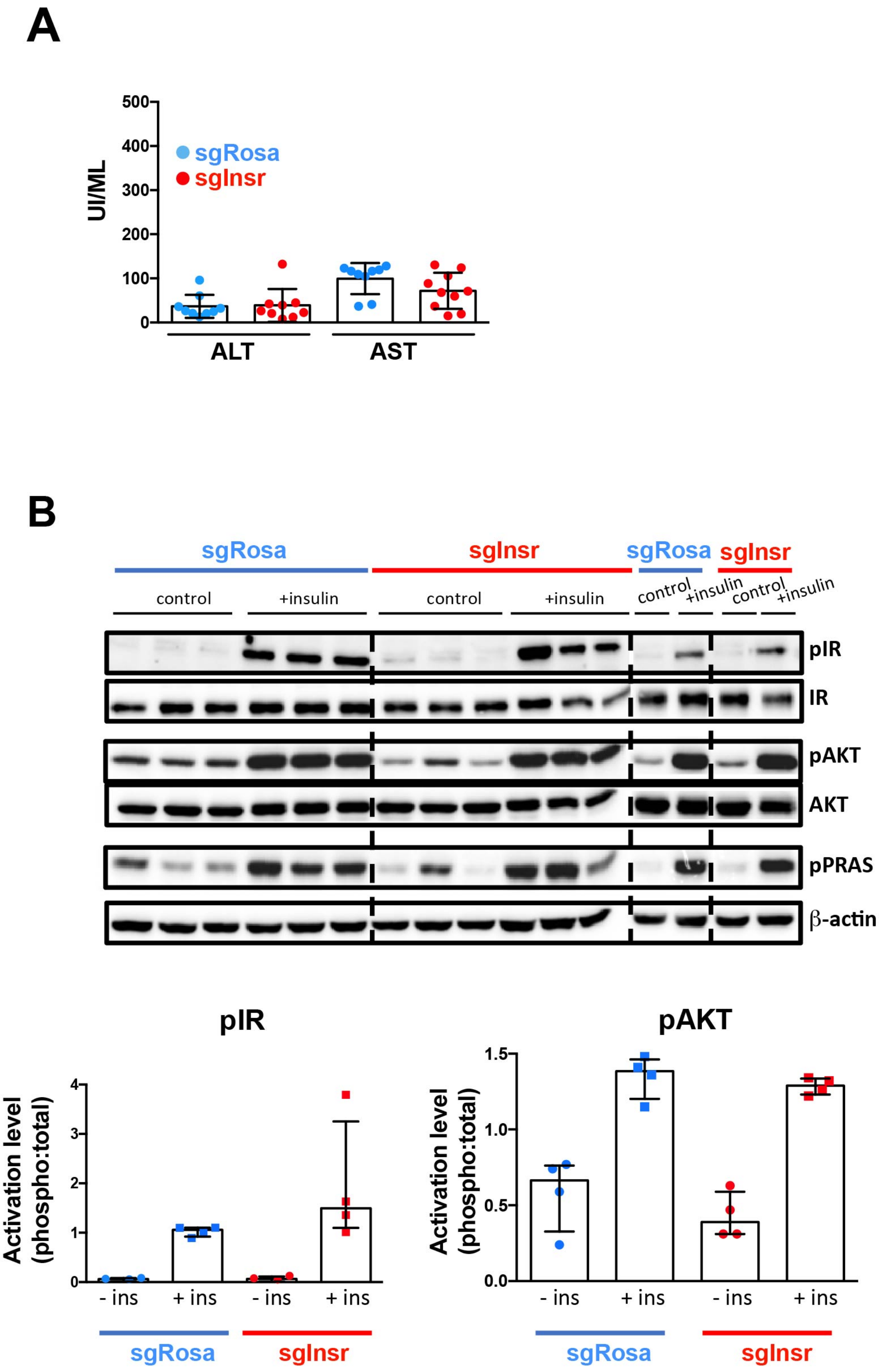
A
UI/ML
sgRosa
sgInsr
ALT
AST
B
sgRosa
sgInsr
control
+insulin
pIR
IR
pAKT
AKT
pPRAS
β-actin
pIR
pAKT
pPRAS
Activation level (phospho:total)
Activation level (phospho:β-actin)
- ins
+ ins

### Figure S5

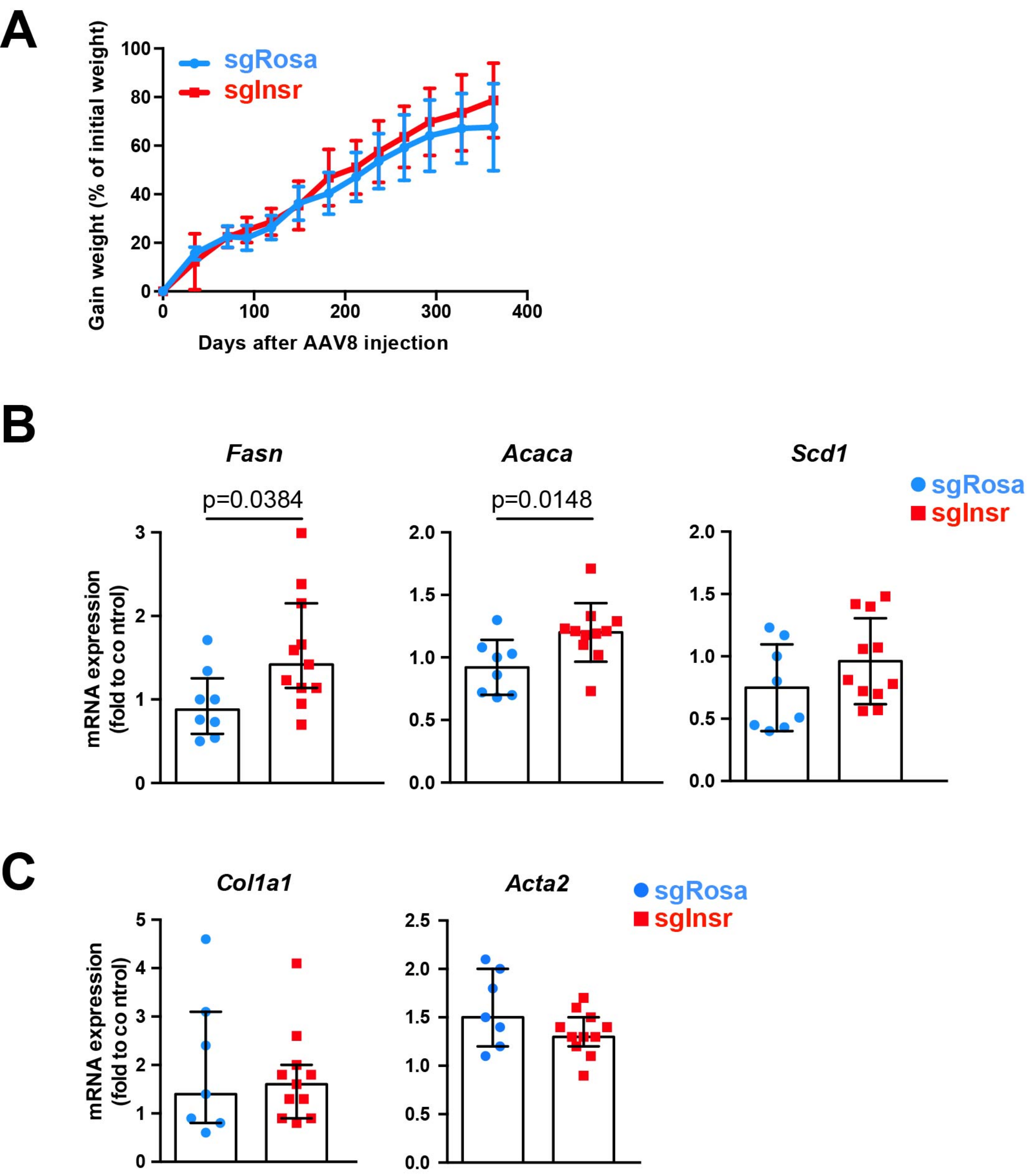
A
Gain weight (% of initial weight)
Days after AAV8 injection
sgRosa
sgInsr
B
Fasn
Acaca
Scd1
p=0.0384
p=0.0148
mRNA expression (fold to co ntrol)
sgRosa
sgInsr
C
Col1a1
Acta2
mRNA expression (fold to co ntrol)
sgRosa
sgInsr

### Figure S6

**A**

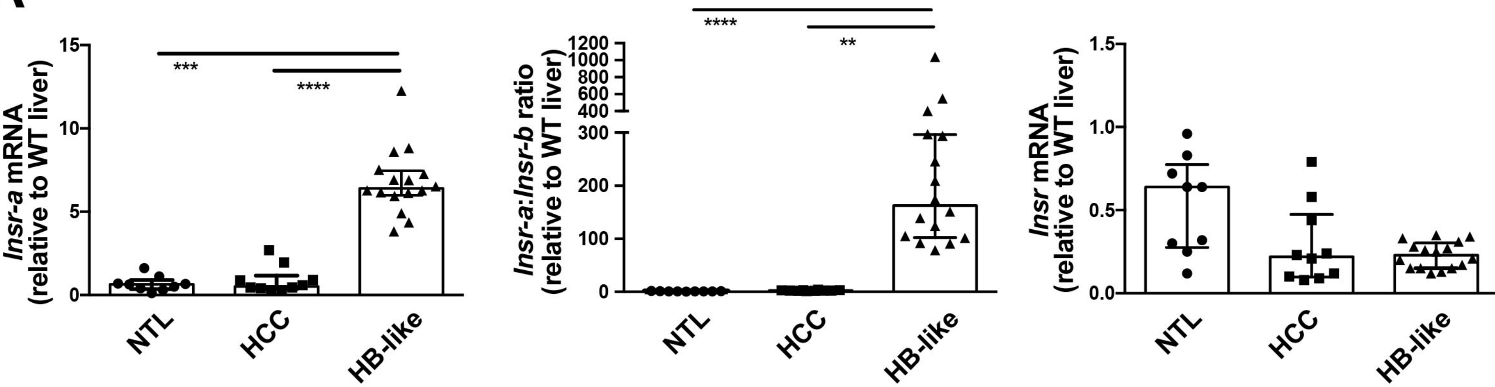

Insr-a mRNA (relative to WT liver)
15
10
5
0
***
****
NTL
HCC
HB-like
Insr-a:Insr-b ratio (relative to WT liver)
1200
1000
800
600
400
300
200
100
0
****
**
NTL
HCC
HB-like
Insr mRNA (relative to WT liver)
1.5
1.0
0.5
0.0
NTL
HCC
HB-like


**B**

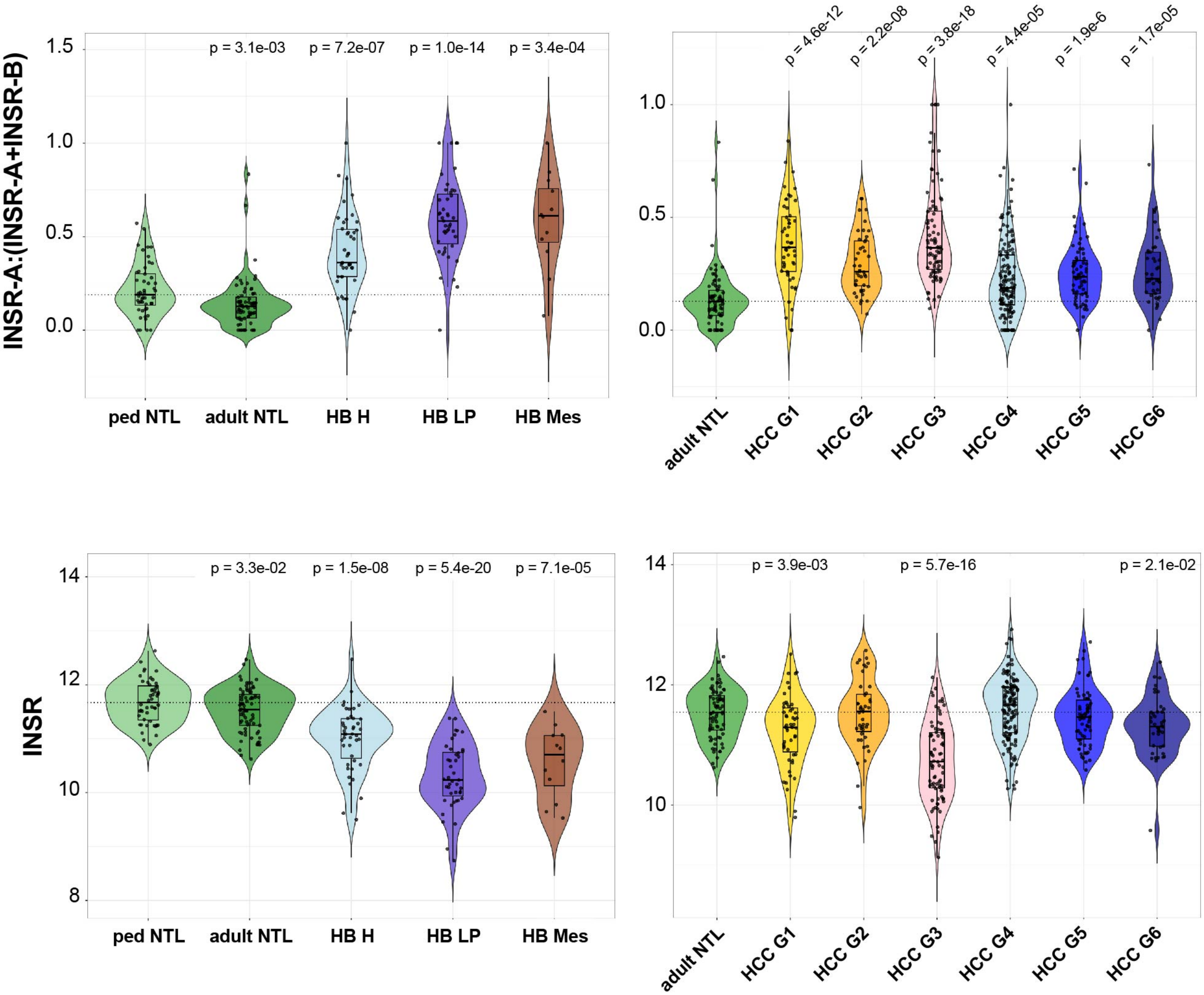

INSR-A:(INSR-A+INSR-B)
1.5
1.0
0.5
0.0
p = 3.1e-03
p = 7.2e-07
p = 1.0e-14
p = 3.4e-04
ped NTL
adult NTL
HB H
HB LP
HB Mes
p = 4.6e-12
p = 2.2e-08
p = 3.8e-18
p = 4.4e-05
p = 1.9e-6
p = 1.7e-05
1.0
0.5
0.0
adult NTL
HCC G1
HCC G2
HCC G3
HCC G4
HCC G5
HCC G6
INSR
14
12
10
8
p = 3.3e-02
p = 1.5e-08
p = 5.4e-20
p = 7.1e-05
ped NTL
adult NTL
HB H
HB LP
HB Mes
14
12
10
p = 3.9e-03
p = 5.7e-16
p = 2.1e-02
adult NTL
HCC G1
HCC G2
HCC G3
HCC G4
HCC G5
HCC G6

Figure S7

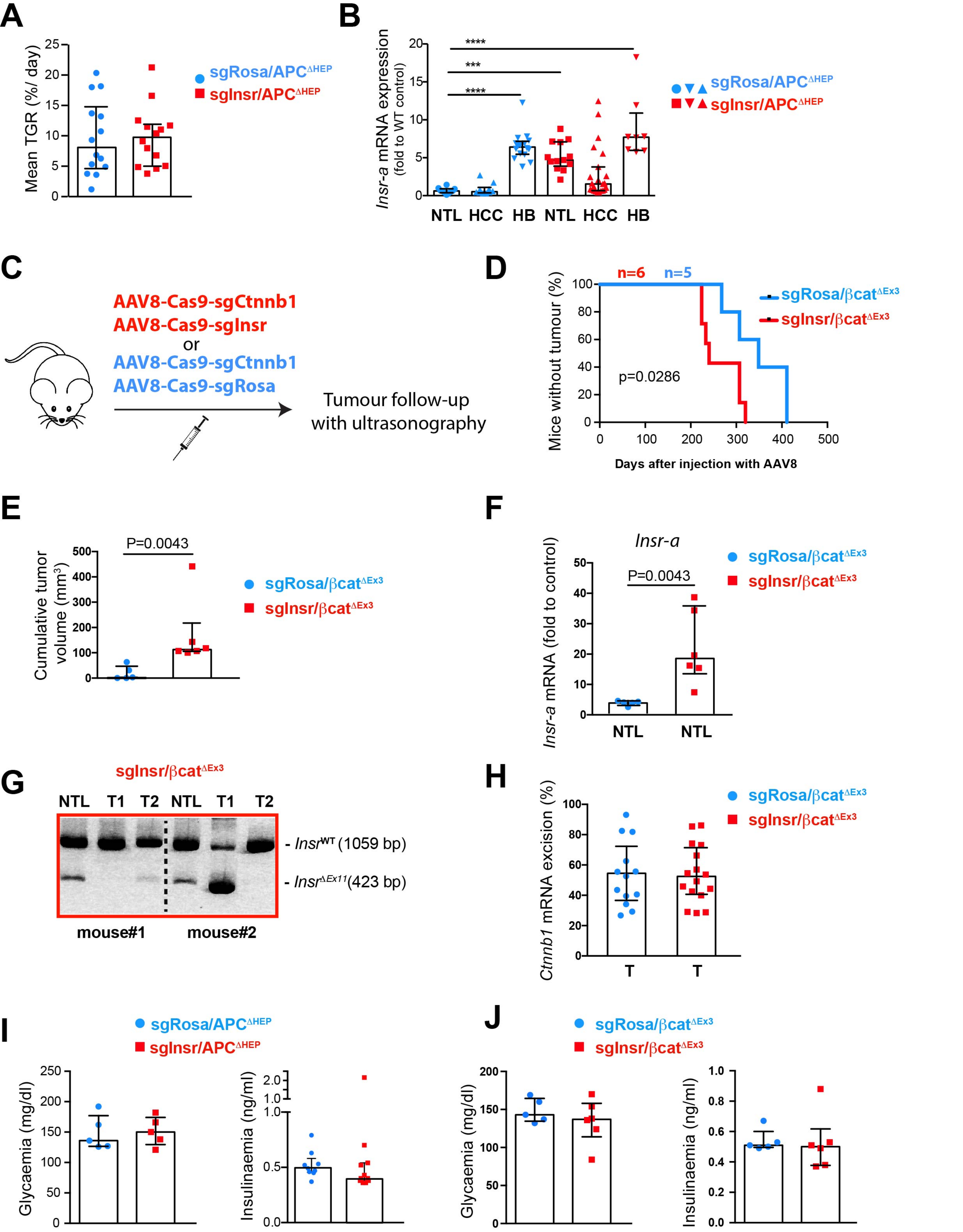

A
Mean TGR (%/ day)
sgRosa/APC$^{\Delta HEP}$
sgInsr/APC$^{\Delta HEP}$
B
Insr-a mRNA expression (fold to WT control)
****
***
****
NTL HCC HB NTL HCC HB
sgRosa/APC$^{\Delta HEP}$
sgInsr/APC$^{\Delta HEP}$
C
AAV8-Cas9-sgCtnnb1
AAV8-Cas9-sgInsr
or
AAV8-Cas9-sgCtnnb1
AAV8-Cas9-sgRosa
Tumour follow-up with ultrasonography
D
n=6 n=5
Mice without tumour (%)
p=0.0286
Days after injection with AAV8
sgRosa/βcat$^{\Delta Ex3}$
sgInsr/βcat$^{\Delta Ex3}$
E
P=0.0043
Cumulative tumor volume (mm$^3$)
sgRosa/βcat$^{\Delta Ex3}$
sgInsr/βcat$^{\Delta Ex3}$
F
Insr-a
P=0.0043
Insr-a mRNA (fold to control)
NTL NTL
sgRosa/βcat$^{\Delta Ex3}$
sgInsr/βcat$^{\Delta Ex3}$
G
sgInsr/βcat$^{\Delta Ex3}$
NTL T1 T2 NTL T1 T2
- Insr$^{WT}$ (1059 bp)
- Insr$^{\Delta Ex11}$(423 bp)
mouse#1 mouse#2
H
Ctnnb1 mRNA excision (%)
T T
sgRosa/βcat$^{\Delta Ex3}$
sgInsr/βcat$^{\Delta Ex3}$
I
sgRosa/APC$^{\Delta HEP}$
sgInsr/APC$^{\Delta HEP}$
Glycaemia (mg/dl)
Insulinaemia (ng/ml)
J
sgRosa/βcat$^{\Delta Ex3}$
sgInsr/βcat$^{\Delta Ex3}$
Glycaemia (mg/dl)
Insulinaemia (ng/ml)

### Figure S8

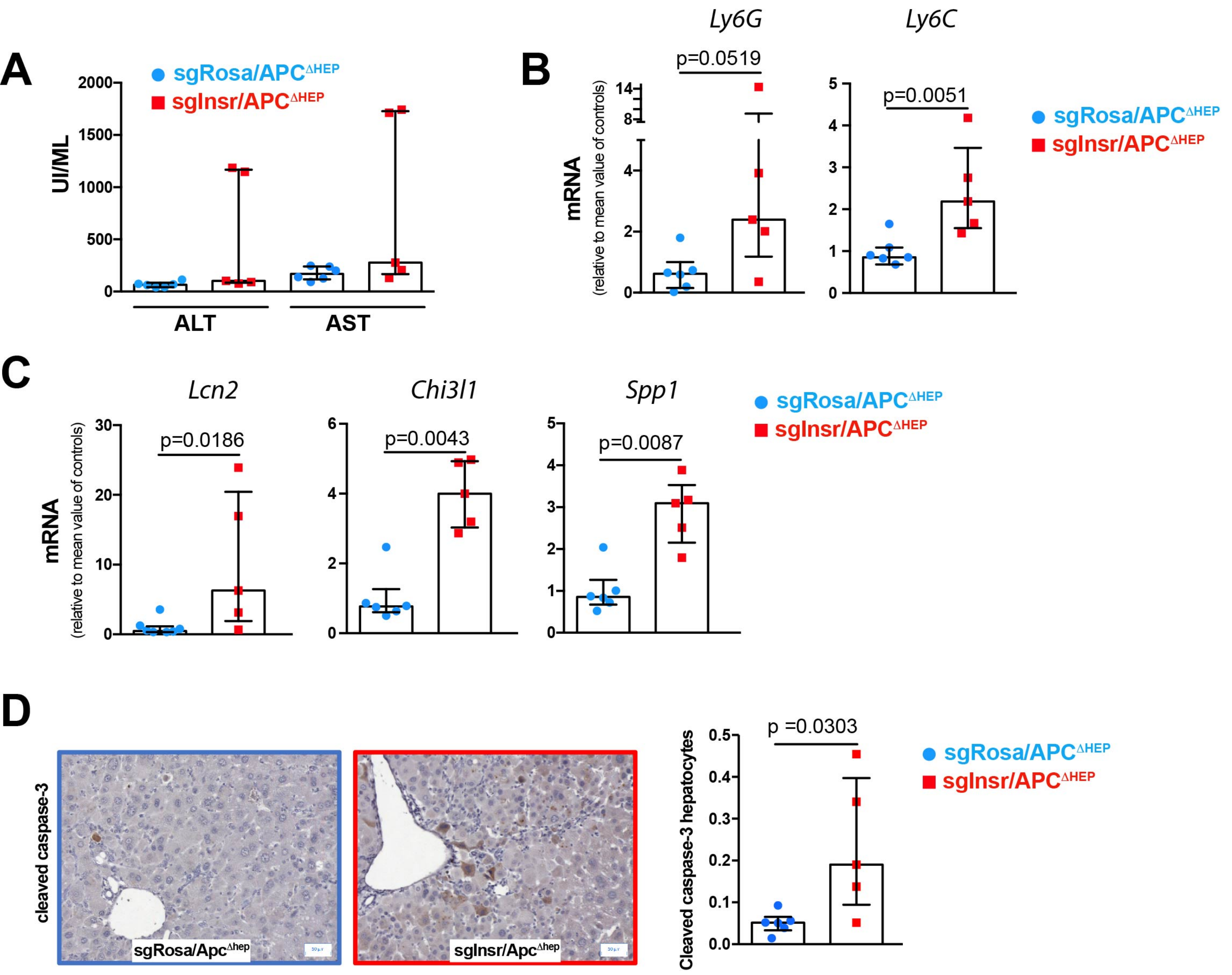


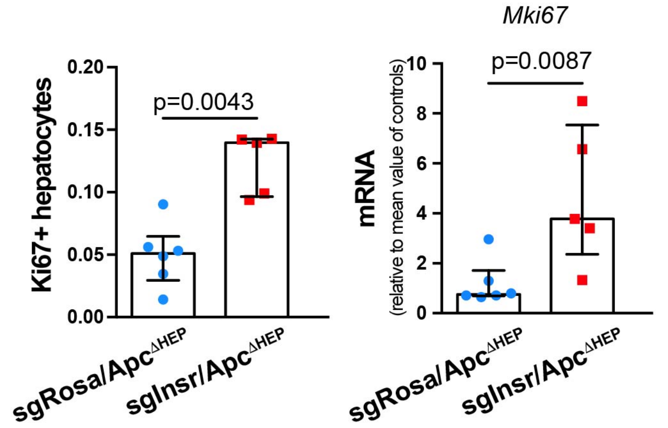


**F**

| | APCΔHep *vs* APC WT | P value |
|---|---|---|
| ***Cxcl2*** | 4.73 | 1.18E-03 |
| ***Chi3l1*** | 3.93 | 7.90E-10 |
| ***Ccl2*** | 3.91 | 2.60E-05 |
| ***Lcn2*** | 2.43 | 1.10E-11 |
| ***Csf1*** | 1.94 | 3.90E-08 |
| ***Mmp9*** | 2.24 | 0.07 |
| ***Mpo*** | 2.24 | ns |
| ***Tnf-α*** | ND | |
| ***Spp1*** | ND | |
| ***Ldlr*** | ND | |

**Legends to supplemental Figures**

**Figure S1: CRISPR/Cas9 tools and design to promote *INSR*/*Insr* exon 11 deletion.**

**A.** Structure diagram of the *pX601*-SaCas9-sgRNA (CMV promoter) and *pX602*-SaCas9-sgRNA (TBG promoter) vectors. **B.** Positions of sgRNAs and primers used for genotyping PCR in the human *INSR* and mouse *Insr* genes.

**Figure S2: CRISPR/Cas9 strategy is efficient in promoting *INSR* exon 11 deletion and favours INSR-A expression in human HCC cells.**

Human PLC/PRF5 cells were stably transfected with two plasmids *pX601*-SaCas9-sgRNA encoding sgRNA hI10-3 and sgRNA hI11-1 directed against *INSR* intron 10 and intron 11 together with a puromycin resistance plasmid and cloned by limiting dilution. **A.** Genotyping PCR at *INSR* locus performed in PLC/PRF5 clones. **B.** Positions of primers used to discriminate INSR-A and INSR-B mRNA by RT-PCR and subsequent gel electrophoresis. **C.** Gel analysis of PCR products encompassing exons 10-12 in *INSR* mRNA. **D.** Evaluation of invasive capacity using Matrigel®-coated Transwells® (n=4). Representative pictures of invaded cells stained with DAPI are shown on the right panel. **E.** Evaluation of clone ability to form spheroids in suspension. Representative pictures are shown. Data are median ± IQR.

**Figure S3: CRISPR/Cas9 strategy is efficient in promoting *INSR* exon 11 deletion in murine hepatocytes *in vitro* and *in vivo*.**

**A.** Murine Hepa1-6 hepatoma cells in primary culture were transiently transfected with plasmids *pX601*-SaCas9-sgRNA encoding sgRNA mI10-1 and sgRNA mI11-1 directed against *Insr* intron 10 and intron 11 or sgRNA against *Rosa* locus, respectively. Genotyping PCR at *Insr* locus are shown. **B.** C57BL/6 mice were injected with increasing doses of recombinant AAV8 encoding SaCas9 under TBG promoter and sgRNA against *Insr* or *Rosa* locus. One month later, livers were harvested and genotyping PCR at *Insr* locus was performed. **C.** Genotyping PCR at *Insr* locus performed using genomic DNA extracted from several organs of a mouse injected with 2.0 x $10^{11}$ vg AAV8-sgInsr. **D.** Representative images of *in situ* mRNA hybridization in the liver. White arrowheads indicate red dots corresponding to *Insr*-a mRNA molecules. Scale bar: 20 $\mu$m.

**Figure S4: Expression of INSR-A in hepatocytes has no impact on liver homeostasis in the short term.**

Mice were injected with recombinant AAV8 encoding saCas9 and sgRNA against *Rosa* (sgRosa) or *Insr* (sgInsr) locus. **A.** Plasma transaminase levels (alanine transaminase (ALT), aspartate transaminase (AST)) determined 11 weeks after injection (n=9-10). **B.** Five weeks after AAV8 injection, mice (n=4 for each group) were injected intraperitoneally with human insulin (1U/kg body weight, 10 min). Saline-injected animals served as controls (n=4 for each group). Whole-liver protein extracts were analyzed by Western blot for intracellular insulin signalling. Band quantification by scanning densitometry is shown below. Data are median ± IQR.

**Figure S5: Expression of INSR-A in hepatocytes leads to tumour development in the long term.**
Mice were injected with recombinant AAV8 encoding saCas9 and sgRNA against *Rosa* (sgRosa, n=10) or *Insr* (sgInsr, n=11) locus and followed by ultrasonography during 13 months before sacrifice. **A.** Weight gain over time. **B, C.** Expression of lipogenic and fibrotic markers in non-tumour livers evaluated by RT-qPCR (n=7-11). Data are median ± IQR except for weight curves (mean± SD).

**Figure S6: Status of *Insr-a/INSR-a* mRNA in murine APC$^{\Delta Hep}$ tumours and human HCC and HB.**
**A.** Analysis of *Insr-a* (*left panel*), *Insr-a*/*Insr-b* ratio (*middle panel*) and total (*right panel*) *Insr* mRNA in APC$^{\Delta Hep}$ HCC and HB-like tumours. Data are median ± IQR. **B.** RNAseq expression data for *INSR-a* and *INSR* in HB (n=98) and HCC (n=427) subclasses normalized to pediatric (n=48) and adult (n=73) non-tumour liver (NTL) tissues, respectively. Ped, pediatric, H, hepatocytic; LP, liver progenitor; Mes, mesenchymal.

**Figure S7: Expression of INSR-A in hepatocytes boosts β-catenin-driven liver carcinogenesis.**
**A.** Mean tumour growth rate (TGR, %/day) evaluated by liver ultrasonography in Apc$^{lox/lox}$ mice consecutively injected with recombinant AAV8 encoding saCas9 and sgRNA against *Rosa26* (sgRosa/APC$^{\Delta Hep}$, n= 14) or *Insr* (sgInsr/APC$^{\Delta Hep}$, n=14) locus and with adenovirus encoding the Cre recombinase. **B.** Expression of *Insr-a* mRNA in HCC and HB tumours (T) relative to adjacent NTL evaluated by RT-qPCR (n=25-28). **C.** Mice were injected with AAV8 encoding sgRNA against *Ctnnb1* together with those targeting *Insr* (sgInsr/βcat$^{\Delta Ex3}$, n=6) or *Rosa* locus (sgRosa/βcat$^{\Delta Ex3}$, n=5) and tumour development was followed by ultrasonography. Schematic protocol is shown. **D.** Tumour appearance over time in sgRosa/βcat$^{\Delta Ex3}$ and sgInsr/βcat$^{\Delta Ex3}$ mice. **E.** Cumulative tumour volumes in sgRosa/βcat$^{\Delta Ex3}$ and sgInsr/βcat$^{\Delta Ex3}$ mice evaluated by ultrasonography 300 days after AAV8 injection (n=5-6). **F.** Expression of *Insr-a* transcripts in nontumour livers (NTL) evaluated by RT-qPCR (n=5-6). **G.** Representative image of *Insr* gene edition evaluated by PCR in nontumour liver (NTL) and tumour (T) tissues. Only tumour T1 from mouse#2 was edited at *Insr* locus. **H.** Expression of excised *Ctnnb1* mRNA in tumours (T) evaluated by RT-qPCR (n=13-16). **I.** Fasting glycaemia (*left*) and insulinaemia (*right*) evaluated in sgRosa/APC$^{\Delta Hep}$ and sgInsr/APC$^{\Delta Hep}$ mice (n=5-9) at sacrifice. **J.** Fasting glycaemia (*left*) and

insulinaemia (*right*) evaluated in sgRosa/β$cat^{\Delta Ex3}$ and sgInsr/β$cat^{\Delta Ex3}$ mice (n=5-6) at sacrifice. Data are median ± IQR.

**Figure S8: Detection of inflammation and apoptosis in sgInsr/$APC^{\Delta hep}$ livers.**

**A.** Plasma transaminase levels (alanine transaminase (ALT), aspartate transaminase (AST)) (n=5-6). **B, C.** Expression of transcripts evaluated by RT-qPCR (n=5-6). **D.** Representative images of cleaved caspase-3 IHC staining from sgRosa/$APC^{\Delta hep}$ and sgInsr/$APC^{\Delta hep}$ livers (*left panel*); Quantification of cleaved caspase-3-positive hepatocytes (*right panel*, n=5-6). **E.** Quantification of Ki67-positive hepatocytes evaluated by IHC (*left panel*) and evaluation of *Mki67* mRNA expression by RT-QPCT (*right panel*) (n=5-6). **F.** Expression of RNAs encoding cytokines/chemokines in $APC^{\Delta hep}$ livers *versus* $APC^{WT}$ livers from RNAseq data. Results are represented as the Log2 fold induction in $APC^{\Delta hep}$ livers *versus* $APC^{WT}$ livers (second column) with their respective p-value. Data are median ± IQR.

**Table S1:** Oligonucleotides used to target *INSR*, *Insr* and *Rosa26* loci with CRISPR/Cas9 strategy

| Species | Targeted genes | Targeted introns | Forward (5'-> 3') | Reverse (5'-> 3') |
|---|---|---|---|---|
| Mouse | *Insr* | 10 | CACCGTGTTACTACAATGAATTACCT | AAACAGGTAATTCATTGTAGTAACA |
| | | 11 | CACCGCCCAGTGAAGGTGTTTAATTG | AAACCAATTAAACACCTTCACTGGG |
| | *Rosa26* | | GATCCGCCTCGGAGTATTTTCCATCGAGG | CTAGGCGGAGCCTCATAAAAGGTAGCTCC |
| Human | *Insr* | 10 | CACCGTCCGTCTAGCAAGTGATGGGA | AAACTCCCATCACTTGCTAGACGGAC |
| | | 11 | CACCGCAGAGTGAAGGCATTGGATTT | AAACAAATCCAATGCCTTCACTCTGC |

**Table S2:** Oligonucleotides used for PCR and qPCR (SybrGreen chemistry)

| Species | Targeted genes | Applications | Forward (5'-> 3') | Reverse (5'-> 3') |
|---|---|---|---|---|
| Mouse | *Insr* | PCR Editing | TTTCCTTCTCCGGATTCCCTTCA | TGCTGAAATTTGGCAACACTGTA |
| | *Gapdh* | PCR Editing | GGCCACGCTAATCTCATTTT | AAGGCGGAGTTACCAGAGGT |
| | *Ctnnb1* | PCR Editing | TTTTGGTGTCGGGGCACATA | CATGGTGCGTACAATGGCAG |
| | *Insr-a* | qPCR | GTTTTTGTCCCCAGGCCAT | TGGCTGTCACA TTCCCCAC |
| | *Insr-b* | qPCR | TTTTGTCCCCAGAAAAACCTCT | GTGGCTGTCACATTCCCCACC |
| | *Insr-tot* | qPCR | TCTTTCTTCAGGAAGCTACATCTG | TGTCCAAGGCATAAAAAGAATAGTT |
| | *Hprt* | qPCR | TCAGTCAACGGGGGACATAA | TGCTTAACCAGGGAAAGCAAA |
| | *Afp* | qPCR | CACACCCGCTTCCCTCAT | TTTTCGTGCAATGCTTTGGA |
| | *Gpc3* | qPCR | CTGTGCTGGAACGGACAAGAAC | GTCAATGATCTGGCTAACCACCG |
| | *Ccl2* | qPCR | TCTGGGCCTGCTGTTCACA | GGATCATCTTGCTGGTGAATGA |
| | *Cxcl2* | qPCR | CCAACCACCAGGCTACAGG | GCGTCACACTCAAGCTCTG |
| | *Csf1* | qPCR | TACAACTGGAAGTGGAGGAGCC AT | AGTCCTGTGTGCCCAGCATAGAAT |
| | *Fasn* | qPCR | TTCCAAGACGAAAATGATGC | AATTGTGGGATCAGGAGAGC |
| | *Acaca* | qPCR | TTACAGGATGGTTTGGCCTTTC | CAAATTCTGCTGGAGAAGCCAC |
| | *Col1a1* | qPCR | TGGAGAGAGCATGACCGATG | TGGACATTAGGCGCAGGAAG |
| | *Acta2* | qPCR | CCACTGAACCCTAAGGCCAAC | AGGGACAGCACAGCCTGAAT |
| | *Ly6g* | qPCR | CTTCTCTGATGGATTTTGCGTTG | AGTAGTGGGGCAGATGGGAAG |
| | *Ly6c* | qPCR | GCAGTGCTACGAGTGCTATGG | ACTGACGGGTCTTTAGTTTCCTT |
| | *Lcn2* | qPCR | ATGTCACCTCCATCCTGGTCAG | GCCACTTGCACATTGTAGCTCTG |
| | *Chi3l1* | qPCR | GCTTTGCCAACATCAGCAGCGA | AGGAGGGTCTTCAGGTTGGTGT |
| | *Spp1* | qPCR | GCTTGGCTTATGGACTGAGGTC | CCTTAGACTCACCGCTCTTCATG |
| | *Mki67* | qPCR | CTGCCTGCGAAGAGAGCATC | AGCTCCACTTCGCCTTTTGG |
| Human | *INSR* | PCR | GCCAGACTTGGAGAAGTGG | CCGCGGGGAGCTCAGA |

**Table S3:** References for TaqMan assays

| Species | Targeted genes | References |
|---|---|---|
| Mouse | *Apc* (exon 10-11) | Mm00545877 |
| | *Apc* (exon 14-15) | Mm01130462 |
| | *Tnfa* | Mm00443258 |
| | *Hprt* | Mm03024075 |
| | *Axin2* | Mm00443610 |
| | *Glul* | Mm00725701 |
| | *Scd1* | Mm01197142 |
